# A dual-domain chitinase mechanism enables marine bacteria to sense and localize sparse crystalline chitin in the ocean

**DOI:** 10.64898/2025.12.08.690172

**Authors:** Hinata Tanimoto, Mikiko Tsudome, Mikako Tachioka, Atsushi Baba, Masayuki Miyazaki, Takayuki Uchihashi, Ryota Iino, Shigeru Deguchi, Akihiko Nakamura

## Abstract

Chitin is one of the most abundant biopolymers in the ocean, and its remineralisation underpins major pathways of marine carbon cycling. A central challenge for chitinolytic bacteria is to locate and remain associated with sparsely distributed crystalline chitin particles. Here, we show that the GH18 chitinase VpChi1 from *Vibrio parahaemolyticus* possesses a dual-domain architecture that enhances particle encounter and retention. Full-length VpChi1 displayed much higher catalytic efficiency than a CBD-truncated mutant at low chitin concentrations, whereas substrate inhibition occurred at higher concentrations. Single-molecule fluorescence imaging demonstrated that the additional C-terminal chitin-binding domain (CBD) enables selective binding to the hydrophobic surface of crystalline chitin and substantially prolongs enzyme residence time. Kinetic simulations using experimentally determined dissociation rates showed that this extended residence arises from CBD-assisted rebinding of the catalytic domain. High-speed atomic force microscopy confirmed that the CBD does not alter stepping velocity or travel distance along individual crystals, indicating that substrate inhibition likely results from CBD engagement with neighbouring particles rather than impaired motility. Screening of seawater and marine sediments across multiple depths identified *Vibrio* and *Pseudoalteromonas* species as dominant chitin degraders, many encoding GH18 chitinases with dual CBDs, whereas terrestrial bacteria generally lacked this architecture. We propose that this two-domain configuration confers an ecological advantage by increasing enzyme retention on particulate chitin and enabling sustained production of chitobiose, a potent chemotactic signal for marine bacteria. These findings reveal a mechanistic basis for how bacteria sense and exploit particulate chitin and highlight an adaptive strategy for resource acquisition in the ocean.

## Introduction

Chitin, a polymer of N-acetylglucosamine, is a major structural component of arthropod exoskeletons and fungal cell walls. Owing to the exceptional stability of its crystalline form—which does not decompose below 320°C under 25 MPa [1], chitin persists extensively in both terrestrial and marine ecosystems [2,3]. In the ocean, copepods dominate the zooplankton community and are key drivers of particulate organic matter export. Their moults, carcasses and associated faecal pellets contribute substantially to marine snow sinking through the water column. It is estimated that more than 1 × 10^6^ tonnes of chitin are produced annually by copepods alone, the majority of which is remineralised by chitin-degrading bacteria [4].

A central ecological challenge for marine chitinolytic bacteria is the initial encounter with insoluble chitin particles dispersed throughout the water column. Many bacteria exhibit chemotaxis towards chitin-derived oligosaccharides, a behaviour predicted to enhance retention near marine snow particles and increase encounter rates [5]. In *Vibrio fischeri*, gradients of chitin degradation products guide migration towards the squid light organ [6]. Chemotactic sensing relies on methyl-accepting chemotaxis proteins whose periplasmic domains detect soluble ligands and regulate motility via the CheA–CheY pathway [7]. These mechanisms highlight the importance of soluble chitin derivatives as ecological cues for particle localisation.

Through extracellular secretion of chitinases, marine chitinolytic bacteria convert recalcitrant insoluble chitin into soluble oligosaccharides, thereby fuelling microbial heterotrophy and shaping carbon cycling. Bacterial chitinases are classified into glycoside hydrolase families (GH) 18 and 19 in the Carbohydrate Active enZymes database [8], and their expression is tightly coupled to nutrient status. *Vibrio furnissii* produces chitinases under starvation, indicating that chitin is a major carbon and nitrogen source [4]. *Vibrio parahaemolyticus*, three GH18 chitinases (VP2338, VPA0055 and VPA1177) and one GH19 enzyme (VP0619) respond to chitin-derived molecules. The secreted deacetylase VP2638 generates *N*-acetyl-*β*-D-glucosaminyl-(1,4)-D-glucosamine (GlcNAc–GlcN), which triggers more than 80-fold induction of the GH18 chitinase gene VPA0055 (VpChi1) [9]. In *Vibrio harveyi*, chitobiose binding to the periplasmic chitooligosaccharide-binding protein VhCBP induces its dissociation from the sensor ChiS, activating downstream chitinase expression [10,11]. Although the regulatory networks governing chitin utilisation are increasingly well understood, the molecular mechanistic basis by which bacteria generate the very signals required for particle detection remains insufficiently explored.

Despite the ecological prominence of chitin in marine systems, most mechanistic insights into chitinase function derive from terrestrial enzymes such as SmChiA and SmChiB from *Serratia marcescens*. These GH18 chitinases comprise a catalytic domain (CD) and a chitin-binding domain (CBD) [12] , and both possess aromatic-lined surfaces that mediate adsorption to crystalline substrates [13]. They hydrolyse chitin processively while moving directionally along the polymer chain, producing diacetylchitobiose (GlcNAc2); SmChiA progresses from the reducing to the non-reducing end, whereas SmChiB moves in the opposite direction [14]. Such single-molecule and biophysical approaches provide powerful tools for examining how marine chitinases interact with natural crystalline substrates, and how these interactions may shape microbial foraging strategies in the ocean.

Here, we investigate the functional contribution of the additional C-terminal CBD in the GH18 chitinase VpChi1, a major chitinase in *V. parahaemolyticus*. Using single-molecule fluorescence imaging, high-speed atomic force microscopy (HS-AFM) and X-ray crystallography, we compare the behaviour of full-length VpChi1 with that of a CBD-truncated variant. We further identify major chitin-degrading bacteria across diverse marine environments and show that GH18 chitinases with dual-CBD architecture are widespread among marine taxa but largely absent from terrestrial lineages. Together, these findings reveal how domain architecture modulates enzyme–particle interactions and provide mechanistic insight into how marine bacteria sense, localise and exploit sparsely distributed crystalline chitin in the ocean.

## Materials and Methods

### Construction of the expression plasmids for VpChi1, VpChi1-truncated and 2ndCBD

The *VpChi1* gene (GenBank: AB299855.1) was codon-optimised for *Escherichia coli* K-12, synthesised with NdeI and NotI sites, digested with the same enzymes (New England Biolabs) and ligated into pET27b (Merck Millipore). VpChi1-truncated was generated by removing the signal peptide (Ile2–Ala21), introducing an N-terminal His₆-tag and TEV site (HHHHHHENLYFQG), and deleting the C-terminal FN3 and CBD regions (Val592–Asn848). PCR fragments and the pET27b backbone were assembled using NEBuilder HiFi DNA Assembly (New England Biolabs).

For fluorescent labelling, the A352C mutation was introduced into both VpChi1 and VpChi1-truncated. The construct encoding the isolated 2ndCBD was produced by PCR, replacing the region Ile2–Val591 of VpChi1 with a His₆-tag, TEV site and a cysteine residue (HHHHHHENLYFQGC).

All plasmids were transformed into *E. coli* Tuner (DE3) (Novagen) by electroporation. Transformants were selected on LB agar containing 50 μg mL^-1^ kanamycin after a 1 h recovery at 37 °C in SOC medium. Single colonies were cultured in 10 mL LB–kanamycin (50 μg mL^-1^) at 37 °C, 300 rpm for 14 h. Plasmids were purified using a QIAprep Spin Miniprep Kit (Qiagen), and sequences were verified by Eurofins Genomics and compared with designed constructs using APE.

### Protein expression

(i) *E. coli* Tuner (DE3) was electroporated with sequence-verified plasmids and recovered for 1 h at 37 °C in SOC medium before plating on LB agar containing 50 μg mL^-1^ kanamycin. Single colonies were grown overnight in 10 mL LB-kanamycin and used to inoculate 700 mL LB–kanamycin (25 μg mL^-1^). Cultures were incubated at 37 °C, 130 rpm for 3 h, chilled on ice for 30 min and induced with 0.5 mM IPTG. Expression proceeded at 20 °C for 14 h before harvesting cells by centrifugation (3,000 g, 20 min).

### Purification of VpChi1 and VpChi1-A352C

Culture supernatants were clarified and concentrated 50-fold using a Minimate TFF system (10 kDa MWCO; Pall). Ammonium sulphate was added to 80% saturation and the suspension was incubated at 4 °C for 16 h. Precipitated protein was collected (20,000 g, 20 min) and resuspended in 20 mM Tris-HCl pH 8 containing 1 M ammonium sulphate.

Samples were applied to a Toyopearl Phenyl-650S column (Tosoh), washed with the same buffer and eluted over a 1-0 M ammonium sulphate gradient. Active fractions were concentrated and desalted (Vivaspin 20, 30 kDa MWCO) and further purified by anion exchange on Toyopearl DEAE-650S in 20 mM Tris-HCl pH 8 using a 0-500 mM NaCl gradient. Final polishing was performed by gel filtration on a Superdex 200 Increase 10/300 column (Cytiva) equilibrated in 20 mM sodium phosphate pH 6. Protein concentrations were determined by A(280) using ε = 162,920 M^-1^ cm^-1^.

### Purification of VpChi1-truncated, VpChi1-truncated-A352C and 2ndCBD

Cell pellets were resuspended in 50 mM sodium phosphate pH 7 with 100 mM NaCl, lysed by sonication on ice and clarified (20,000 g, 20 min). Supernatants were applied to Ni-NTA agarose (Qiagen), washed with the same buffer and eluted with 50 and 100 mM imidazole. Proteins were concentrated (Vivaspin 20; 30 kDa MWCO for truncated constructs, 10 kDa for 2ndCBD), desalted using a NAP-10 column and digested with TEV protease at 16 °C for 14 h.

TEV-digested samples were passed over Ni-NTA again to remove cleaved tags. VpChi1-truncated and VpChi1-truncated-A352C were purified by gel filtration on a Superdex 200 Increase 10/300 column in 20 mM sodium phosphate pH 6, and concentrations were determined using ε = 113,440 M^-1^ cm^-1^.

VpChi1-2ndCBD-addC591 was purified on a Superdex 75 Increase 10/300 column in 20 mM sodium phosphate pH 7, followed by anion exchange on Toyopearl DEAE-650S using a stepped 0-200 mM NaCl gradient. Concentrations were determined using ε = 49,480 M^-1^ cm^-1^.

### Cy3 labeling of enzymes

Enzymes were reduced with 10 mM DTT at 25 °C for 1 h and desalted using a NAP-5 column (Cytiva) into 20 mM sodium phosphate pH 7 containing 100 mM NaCl. Cy3–maleimide (Cytiva) was added at a 1:1 molar ratio and the reaction was incubated for 1 h at 25 °C. The mixture was concentrated to 100 μL (Vivaspin 500) and unreacted dye was removed using a second NAP-5 column. Labelling efficiency was calculated from absorbance at 280 and 550 nm using extinction coefficients of 12,000 M^-1^ cm^-1^ (280 nm) and 150,000 M^-1^ cm^-1^ (550 nm) for Cy3.

### Crystallization and X-ray crystal structure analysis of VpChi1-truncated

VpChi1-truncated (20 mg mL^-1^) was mixed 1:1 with reservoir solution (10% PEG 8000, 100 mM Ca(OAc)2, 100 mM sodium cacodylate pH 6.5) and equilibrated at 20 °C in a sitting-drop plate. Crystals were transferred to reservoir solution containing 30% (v/v) glycerol and flash-cooled.

Diffraction data (λ = 1.54 Å) were collected at 93 K using an iMSx system with 1° oscillation and 20 s exposure. Seven data sets were recorded based on pre-analysis. Data were processed with CrysAlisPro, and phases were obtained by molecular replacement in Phaser using a SWISS-MODEL homology model. The structure was refined using Phenix.refine and manually adjusted in Coot.

### Biochemical assay of crystalline chitin degradation

Crystalline chitin was purified from *Lamellibrachia satsuma* tubes as described previously [15]. Enzyme solutions (100 μL, 200 nM) in 100 mM sodium phosphate pH 6.0 were mixed with crystalline chitin (0-4 mg mL^-1^) and incubated for 15 min at 25 °C. Reactions were terminated with 300 μL of 500 mM sodium carbonate containing 1.5 mM potassium ferricyanide, centrifuged (15,000 g, 5 min, 4 °C) and supernatants were heated at 98 °C for 10 min. Absorbance at 420 nm was compared with chitobiose standards. Activities in artificial seawater were assessed similarly, replacing buffer with two-fold concentrated Daigo Artificial Seawater SP.

### Binding and dissociation rate constant analysis by TIRFM

Coverslips were immersed in 10 M KOH overnight at room temperature, rinsed thoroughly with ultrapure water, and coated with crystalline chitin by spin-casting 50 μL of a 0.01 mg mL^-1^ suspension at 3,000 rpm. The coated surface was mounted on the microscope stage, and regions containing chitin fibrils were selected by bright-field imaging.

Enzyme solutions (25 pM for VpChi1 and VpChi1-truncated; 40 pM for 2ndCBD) in 50 mM sodium phosphate buffer pH 6.0 were applied (20 μL) and observed under 532-nm excitation (0.14 μW μm^-2^ for VpChi1/VpChi1-truncated; 0.28 μW μm^-2^ for 2ndCBD). Imaging rates were 4 fps for VpChi1-A352C-Cy3 and VpChi1-truncated-A352C-Cy3, and 10 fps for VpChi1-2ndCBD-addC591.

Following single-molecule recording, 20 μL of 10 nM enzyme was added to visualise the chitin fibrils fully. A “stained chitin” reference image was generated by averaging 100 frames. Molecules were verified as chitin-bound by overlaying the single-molecule trajectories with this reference.

Binding durations and the number of binding events within a 40-s window were quantified. Dissociation rate constants (koff^fast^ and *k*off^slow^) were obtained by fitting the binding-time distributions with a sum of two exponential decay functions. Binding rate constants (*k*on) were calculated from binding frequencies normalised by observation time, chitin length, and enzyme concentration. The *k*on distributions were fitted with a sum of Gaussian functions, and the lowest-rate peak was taken to represent binding to a single chitin crystal.

### Simulation of binding-time distributions of VpChi1

Binding-time distributions for VpChi1-truncated and the 2^nd^-CBD were first generated using the Python script shown in Supplementary Fig. S7. Random numbers (0-1) were converted to binding times based on the experimentally determined *k*off^slow^ values. Five hundred binding events were simulated, and the resulting distributions were fitted with a single-exponential decay function to confirm that the simulated *k*off values matched the experimental measurements. For VpChi1, binding and dissociation times for each of the two domains were generated from their *k*off^slow^ values, and the binding time of the whole enzyme was taken as the time until both domains had dissociated (Supplementary Fig. S8). Binding-time distributions (n = 500) were then produced for a range of rebinding rate constants (*k*on = 0.25–10 s^-1^). Each distribution was fitted with a single-exponential decay function to obtain the simulated *k*off, and the relationship between *k*off and the rebinding *k*on was fitted using the equation derived from the sequential dissociation model of VpChi1 (Supplementary Fig. S3).

To examine whether *k*on^CD^ and *k*on^CBD^ could be considered equivalent, additional simulations were performed with *k*on^CD^ values of 0.25–10 s^-1^ and *k*on^CBD^ values ranging from one-tenth to tenfold of *k*on^CD^ (Supplementary Fig. S4). Each simulated distribution was fitted with both single- and double-exponential decay functions, and the difference in R^2^ values was used as an indicator of deviation from single-exponential behaviour. Because the two fitting functions become identical for a true single-exponential, the R^2^ difference approaches zero.

Simulated binding-time distributions were considered consistent with the experimental data if the following criteria were met:

i. simulated *k*off fell within 0.049 ± 0.005 s^-1^,
ii. R^2^ for the single-exponential fit exceeded 0.95, and
iii. the difference in R² values between the single- and double-exponential fits was ≤ 0.015.

### HS-AFM observation of VpChi1 and VpChi1-truncated

High-speed atomic force microscopy (HS-AFM) was used to analyse the movement of VpChi1 and VpChi1-truncated, following the procedure previously established for SmChiA [16]. Crystalline chitin was immobilised on a fluorosurf-coated mica substrate, and unbound fibrils were removed by washing. Chitin crystals were located in 78 μL of 50 mM sodium phosphate buffer (pH 6.0), after which 2 μL of 20 μM VpChi1 or VpChi1-truncated was added to initiate the observation. Molecular movement along the crystalline chitin surface was recorded at 5 fps at 25 °C.

### Screening of chitin-degrading bacteria from the sea

Nanofibrous chitin gel was prepared following previously described protocols [17,18]. Briefly, 1.5 g Chitosan 10 powder (FUJIFILM Wako) was dissolved in 54 mL water containing 6 mL acetic acid, diluted with 240 mL methanol, and N-acetylated by adding 1.8 mL acetic anhydride with gentle stirring. The mixture (60 mL per dish) was dispensed into silane-coated glass Petri dishes (15 cm diameter), sealed, and left overnight at room temperature to allow formation of a porous nanofibrous chitin network. The resulting gels were thoroughly washed, autoclaved, transferred to sterile dishes and stored at 4 °C until use.

Sampling operations and isolation of the bacterial strains were performed by a similar method as described in the previous report [19]. *In situ* sampling was performed by exposing nanofibrous chitin gel to seawater in plastic mesh bags fixed to the exterior of the unmanned remotely operated vehicle Hyper-Dolphin (dive nos. 682, 683, and 686 during cruise NT07-09, Jun 2007) or the manned submersible Shinkai 6500 (dive nos. 1029–1035 during cruise YK07-14, Sep-Oct 2007). Sampling conditions are summarised in Table S2. The recovered plates were crushed into fine particles and dispersed in artificial seawater (Marine Art SF-1; Senju Pharmaceutical, Osaka, Japan). Then the dispersions were spread on fresh chitin plates containing artificial seawater and incubated at 20 °C. Isolations were performed by selecting the pits formed on the surfaces of chitin plates and spreading them repeatedly on fresh chitin plates until pure cultures were obtained. The 16S rRNA genes of the isolates were amplified through PCR using the domain bacteria-specific primers 27f (5′ AGAGTTTGATCCTGGCTCAG 3′) and the universal primer 1492r (5′ GGTTACCTTGTTACGACTT 3′), and the PCR product was sequenced using the primer 520r (5′ ACCGCGGCTGCTGGC 3′) as described previously [20].

### Sequence analysis of chitinases

A total of 270 GH18 chitinase sequences were retrieved using the NCBI BLASTp suite with SmChiA or VpChi1 as queries. Redundant sequences from the same organism were removed, and collection was continued until both queries returned overlapping hits. Signal peptides were predicted using SignalP 5.0 (https://services.healthtech.dtu.dk/service.php?SignalP-5.0) and removed prior to alignment. Sequence alignment was performed with the COBALT server (https://www.ncbi.nlm.nih.gov/tools/cobalt/), and the phylogenetic tree was visualised using FigTree v1.4.

## Results

### Structural features of VpChi1

The major chitinase gene that is strongly induced by chitin in *Vibrio parahaemolyticus* (VpChi1) has been identified previously [9], and most *Vibrio* species possess a homologous gene. The molecular mass of this Vibrio chitinase is greater than that of SmChiA [21]. Consequently, the overall architecture of VpChi1 was predicted using ColabFold (Fig. 1A) [22]. The catalytic domains of SmChiA and VpChi1 are highly similar; however, VpChi1 contains two additional fibronectin type III-like (FN3) domains and a C-terminal carbohydrate-binding module of the CBM5 family.

**Fig. 1.**
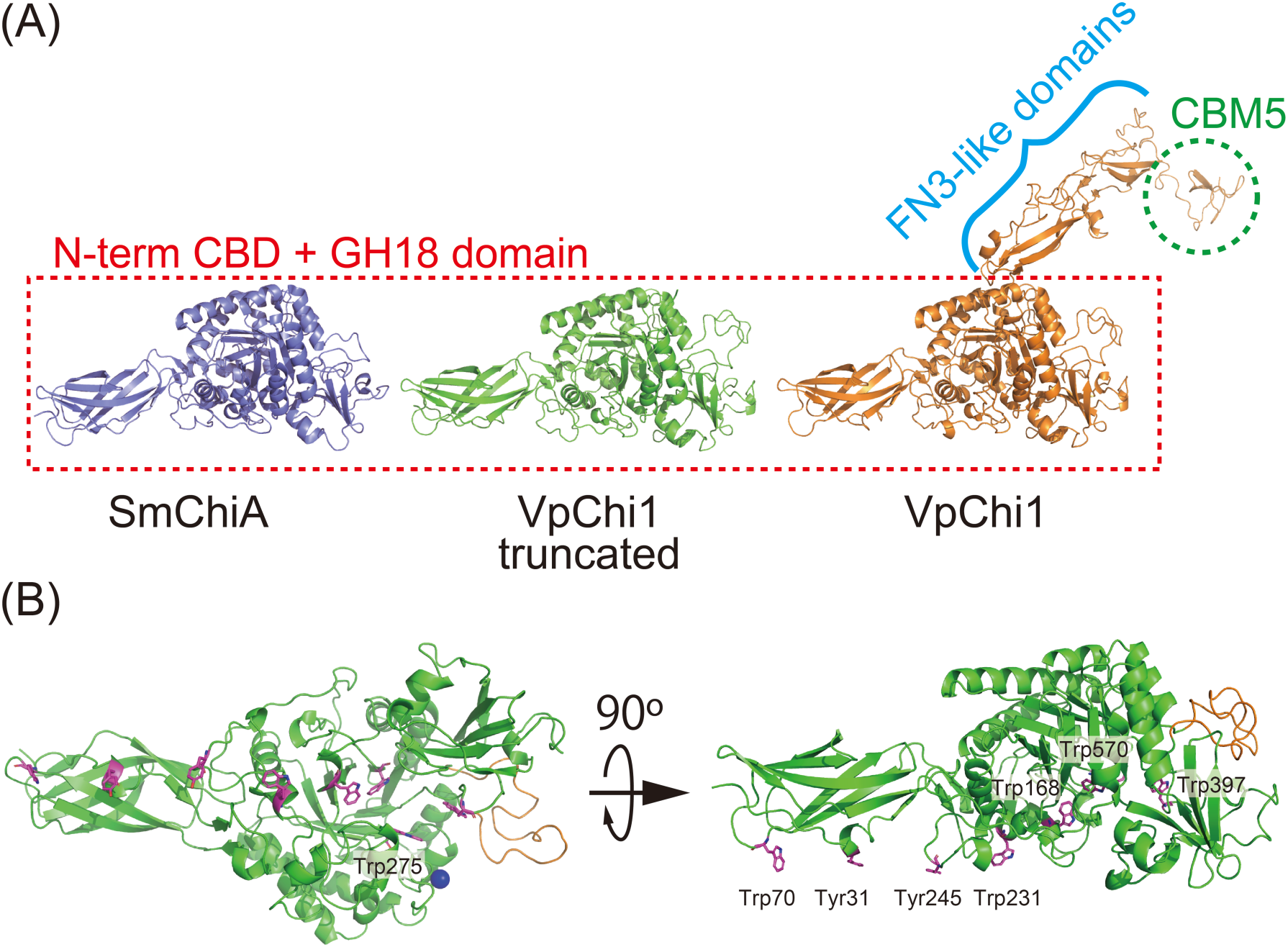
Structure of VpChi1. (A) Predicted domain architectures of SmChiA, VpChi1 and the VpChi1-truncated mutant. All enzymes possess an N-terminal chitin-binding domain (N-term CBD) and a GH18 catalytic domain. VpChi1 additionally contains two fibronectin type III–like (FN3-like) domains and a C-terminal CBM5 chitin-binding domain. Full-length structural models of VpChi1 were generated using ColabFold. (B) X-ray crystal structure of the VpChi1-truncated mutant (PDB ID: 8HRF). VpChi1 exhibits a row of aromatic residues (shown in purple) along the substrate-binding surface, similar to SmChiA, although Tyr31 and Tyr245 replace two of the tryptophan residues found in SmChiA. A calcium ion (shown in blue) is positioned on the molecular surface, and the loop region that is extended relative to SmChiA is highlighted in orange.

The X-ray crystal structure of the catalytic domain of VpChi1 (VpChi1-truncated) was determined at 2.5 Å resolution (Table S1). The catalytic domain adopts an (α/β)₈-barrel fold with an N-terminal chitin-binding domain (Fig. 1B). A flat substrate-binding surface is formed by the aromatic residues Tyr31, Trp70, Trp231 and Tyr245. Additional aromatic residues surrounding the catalytic centre—Trp168, Trp570 and Trp397—and the conserved tryptophan that stabilises the chain-sliding intermediate (Trp275) are also present in VpChi1.

A notable structural distinction is that VpChi1 possesses an extended loop region of 28 residues near the product-binding site, compared with 11 residues in SmChiA. Moreover, Asp367 and Asp369 coordinate a calcium ion in VpChi1, whereas the corresponding positions in SmChiA are occupied by Asp368 and Lys372, which fill the space without metal coordination.

### Effect of additional binding domain to crystalline chitin degradation activity

To investigate the role of the additional binding domain, we compared the biochemical activities of crystalline-chitin degradation between the full-length enzyme (VpChi1) and a mutant lacking the additional FN3 and C-terminal binding domains (VpChi1-truncated) in 50 mM sodium phosphate buffer (pH 6.0). The activity plots of VpChi1-truncated as a function of crystalline chitin concentration were well fitted by the Michaelis–Menten equation, yielding *k*cat and Km values of 4.1 s^-1^ and 0.50 mg mL^-1^, respectively (Fig. 2A). In contrast, VpChi1 displayed maximal activity at low substrate concentrations (0.25 mg mL^-1^), followed by a decrease in activity at higher concentrations, indicative of substrate inhibition (Fig. 2A). Accordingly, its activity data were fitted using a Michaelis–Menten model incorporating a substrate-inhibition constant (Ks) (Fig. 2B). The resulting *k*cat, Km and Ks values were 5.4 s^-1^, 0.055 mg mL^-1^ and 2.1 mg mL^-1^, respectively. These findings indicate that the affinity of the full-length enzyme for crystalline chitin is approximately tenfold higher than that of VpChi1-truncated.

**Fig. 2.**
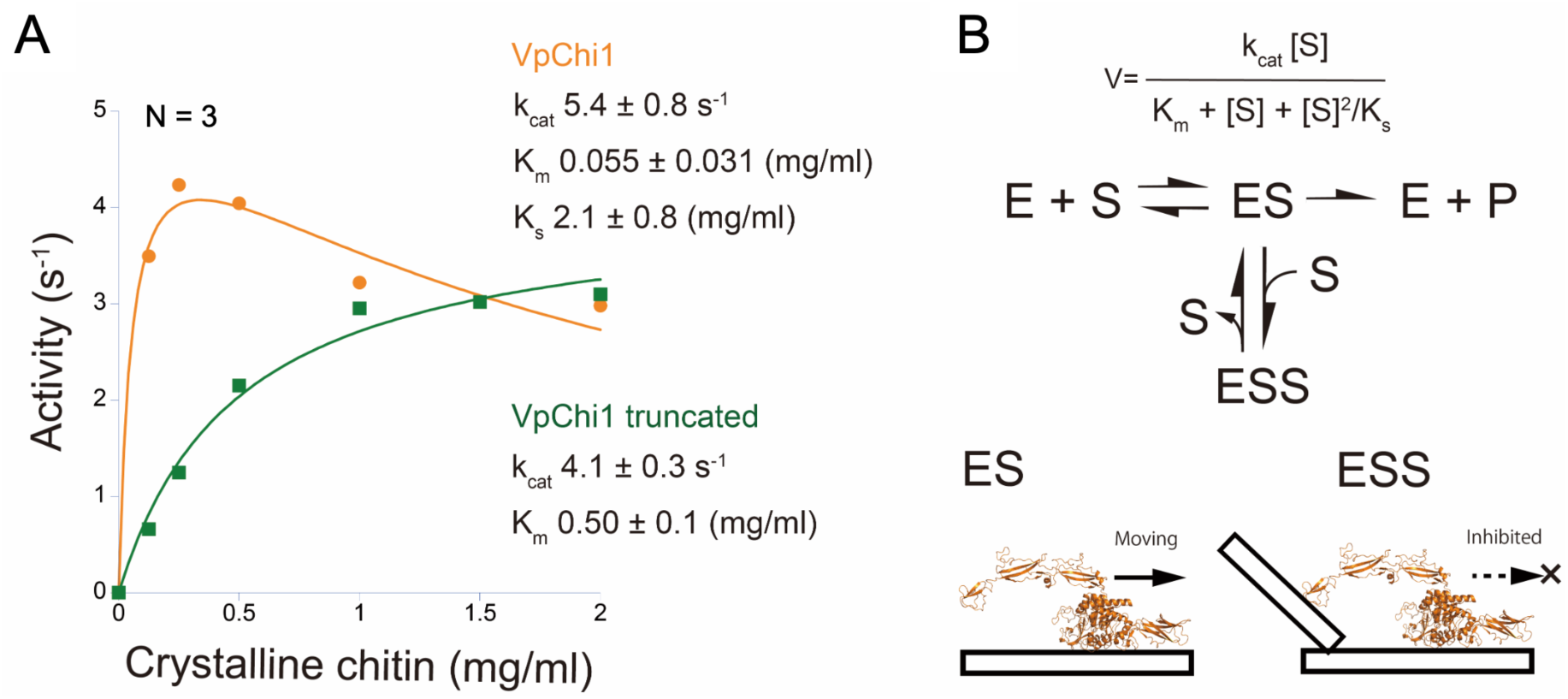
Biochemical activities of VpChi1 and VpChi1-truncated. (A) Activity plots of VpChi1 and VpChi1-truncated as a function of crystalline chitin concentration. Data are presented as mean ± fitting error. (B) Schematic representation of the substrate inhibition mechanism proposed for VpChi1.

We further compared enzyme activities in artificial seawater (pH 7.5). Only VpChi1 exhibited substrate inhibition at high chitin concentrations (Supplementary Fig. S1). Under these conditions, the *k*cat, Km and Ks values of VpChi1 were 1.5 s^-1^, 0.018 mg mL^-1^ and 5.2 mg mL^-1^, respectively, whereas VpChi1-truncated exhibited *k*cat and Km values of 1.3 s^-1^ and 0.42 mg mL^-1^.

At high substrate concentrations, the observed activity of VpChi1 exceeded that predicted by the standard substrate-inhibition model. We therefore evaluated an alternative kinetic model that incorporates hydrolysis by an enzyme simultaneously engaging two chitin crystals (Supplementary Fig. S2A). Using this model, the activity profiles of VpChi1 were fitted more accurately under both buffer and seawater conditions (Supplementary Fig. S2B). However, these results should be regarded as supportive rather than definitive, as the high degrees of freedom in the model introduced substantial fitting uncertainty.

### Experimental analysis of binding and dissociation rate constants

To elucidate why VpChi1 exhibits a smaller Km than VpChi1-truncated, we examined binding and dissociation events on crystalline chitin at the single-molecule level. Free-cysteine mutants of VpChi1, VpChi1-truncated and the isolated C-terminal binding domain (2ndCBD) were prepared and site-specifically labelled with Cy3-maleimide. All enzymes displayed specific binding to crystalline chitin (Supplementary Movies 1–3). The binding rate constant (*k*on) was determined from the number of binding events normalised by observation time, enzyme concentration and the length of the chitin crystal. Strictly, *k*on should scale with substrate surface area; however, because individual crystals are ∼40 nm in width and bundled crystals cannot be resolved by optical microscopy, we statistically estimated *k*on for a single crystal using the smallest peak in each distribution (Fig. 3, upper panels).

**Fig. 3.**
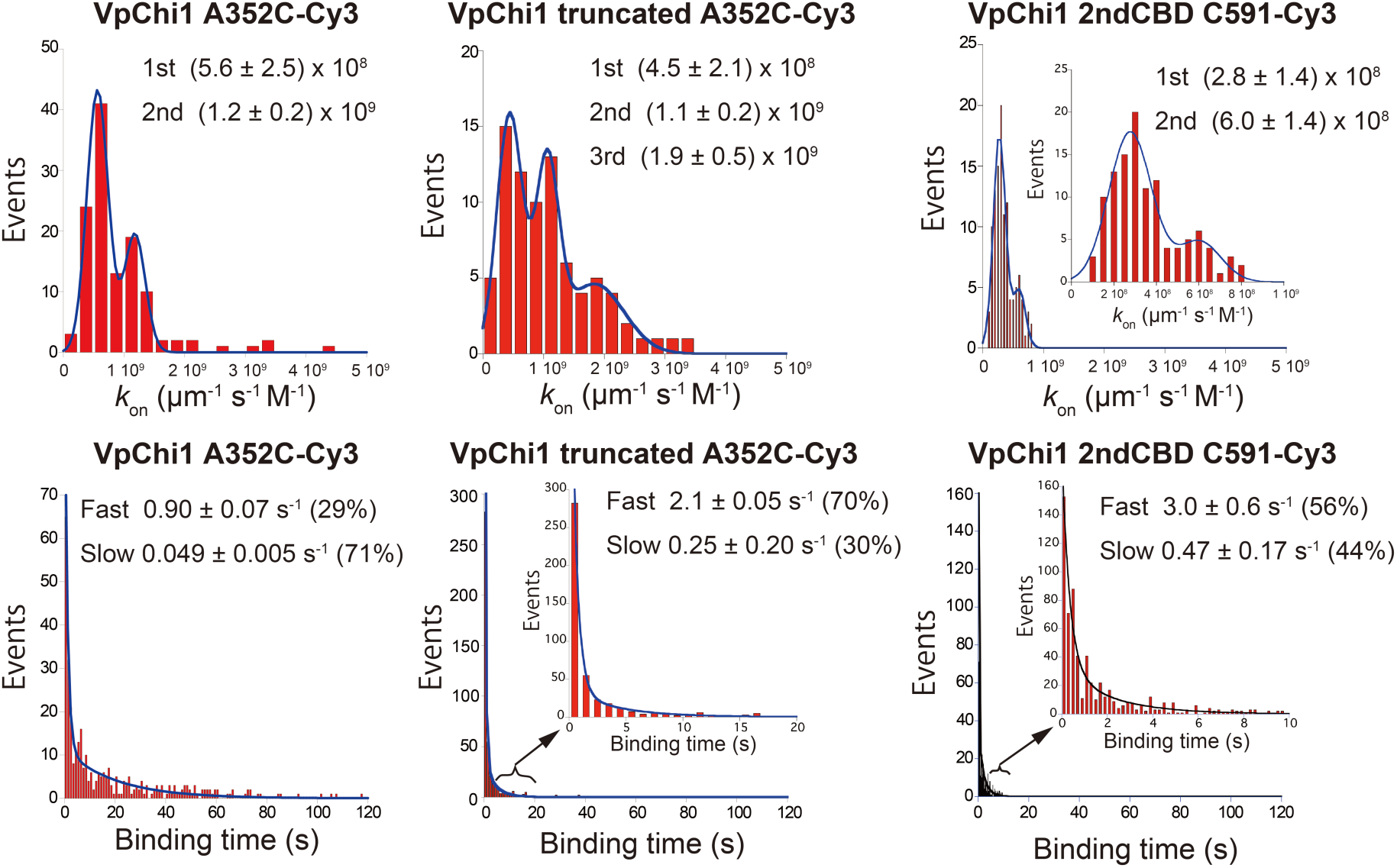
Analysis of binding and dissociation rate constants of VpChi1 and truncated mutant. Distributions of binding rate constants were fitted using a sum of Gaussian functions, whereas binding-time distributions were fitted using a sum of exponential decay functions to resolve fast and slow dissociation states.

VpChi1 showed two major peaks, well fitted by a sum of Gaussian functions, with values of 5.6 × 10⁸ μm^-1^ s^-1^ M^-1^ and 1.2 × 10⁹ μm^-1^ s^-1^ M^-1^. VpChi1-truncated exhibited three peaks at 4.5 × 10⁸ μm^-1^ s^-1^ M^-1^ 1.1 × 10⁹ μm^-1^ s^-1^ M^-1^ and 1.9 × 10⁹ μm^-1^ s^-1^ M^-1^. The *k*on distribution of the 2ndCBD was lower than those of both VpChi1 constructs, with a main peak at 2.8 × 10⁸ μm^-1^ s^-1^ M^-1^ and a secondary peak at 6.0 × 10^8^ μm^-1^ s^-1^ M^-1^.

Dissociation rate constants (*k*off) were obtained from binding-time distributions. All three enzymes exhibited biphasic dissociation kinetics, and the distributions were well fitted by two exponential decay functions yielding *k*off^Fast^ and *k*off^Slow^ (Fig. 3, lower panels). For VpChi1, *k*off^Fast^ and *k*off^Slow^ were 0.90 ± 0.07 s^-1^ and 0.049 ± 0.005 s^-1^, with 29% and 71% of events occurring in the fast and slow states, respectively. VpChi1-truncated predominantly occupied the fast dissociation state (2.1 ± 0.05 s^-1^, 70%), while its slow state (0.25 ± 0.2 s^-1^) represented only 30%. The 2ndCBD displayed the fastest dissociation in both states, with *k*off^Fast^ = 3.0 ± 0.6 s^-1^ and *k*off^Slow^ = 0.47 ± 0.17 s^-1^, and the proportions of fast and slow states were comparable (56% and 44%, respectively).

### Simulation of binding-time distribution for slow dissociation events of VpChi1

The binding-time distribution of slow dissociation events for VpChi1 was simulated using the distributions obtained for VpChi1-truncated and the 2ndCBD, as the slow-dissociation state represents the predominant binding mode and is directly linked to VpChi1 catalytic activity. Dissociation of VpChi1 is assumed to occur when its two domains sequentially detach from the chitin surface; if one domain dissociates while the other remains bound, the dissociated domain can rebind.

We first generated simulated binding-time distributions for VpChi1-truncated and the 2ndCBD using the experimentally determined *k*off^Slow^ values obtained from single-molecule analysis. The simulated *k*off^Slow^ values calculated from 500 dissociation events were 0.24 ± 0.01 s^-1^ for VpChi1-truncated and 0.43 ± 0.02 s^-1^ for the 2ndCBD, closely matching the experimentally measured values (Fig. 4A). For the simulation of VpChi1 binding times, the rebinding rates of the CD and CBD were modelled separately (Fig. 4B and Fig. S3), although they were initially assumed to be identical. Rebinding rate constants (*k*on = 10–0.25 s^-1^) were tested to represent the probability that a dissociated domain rebinds while the other remains attached. The simulated *k*off for VpChi1 decreased progressively as the rebinding *k*on increased (Fig. 4C). The resulting plots were fitted using the equation derived from the sequential dissociation model. When *k*_on_ ^CD^ and *k*_on_ ^CBD^ were fixed to the same value, the estimated dissociation rate constants were *k*₃ = 0.025 ± 0.006 s^-1^ (CD dissociation following CBD dissociation) and *k*₄ = 0.56 ± 0.05 s^-1^ (CBD dissociation following CD dissociation). When the rebinding *k*on was set to 2.3 s^-1^, the simulated distribution was well described by a single-exponential decay and successfully reproduced the experimental value, yielding *k*off^VpChi1^ = 0.049 ± 0.002 s^-1^.

**Fig. 4.**
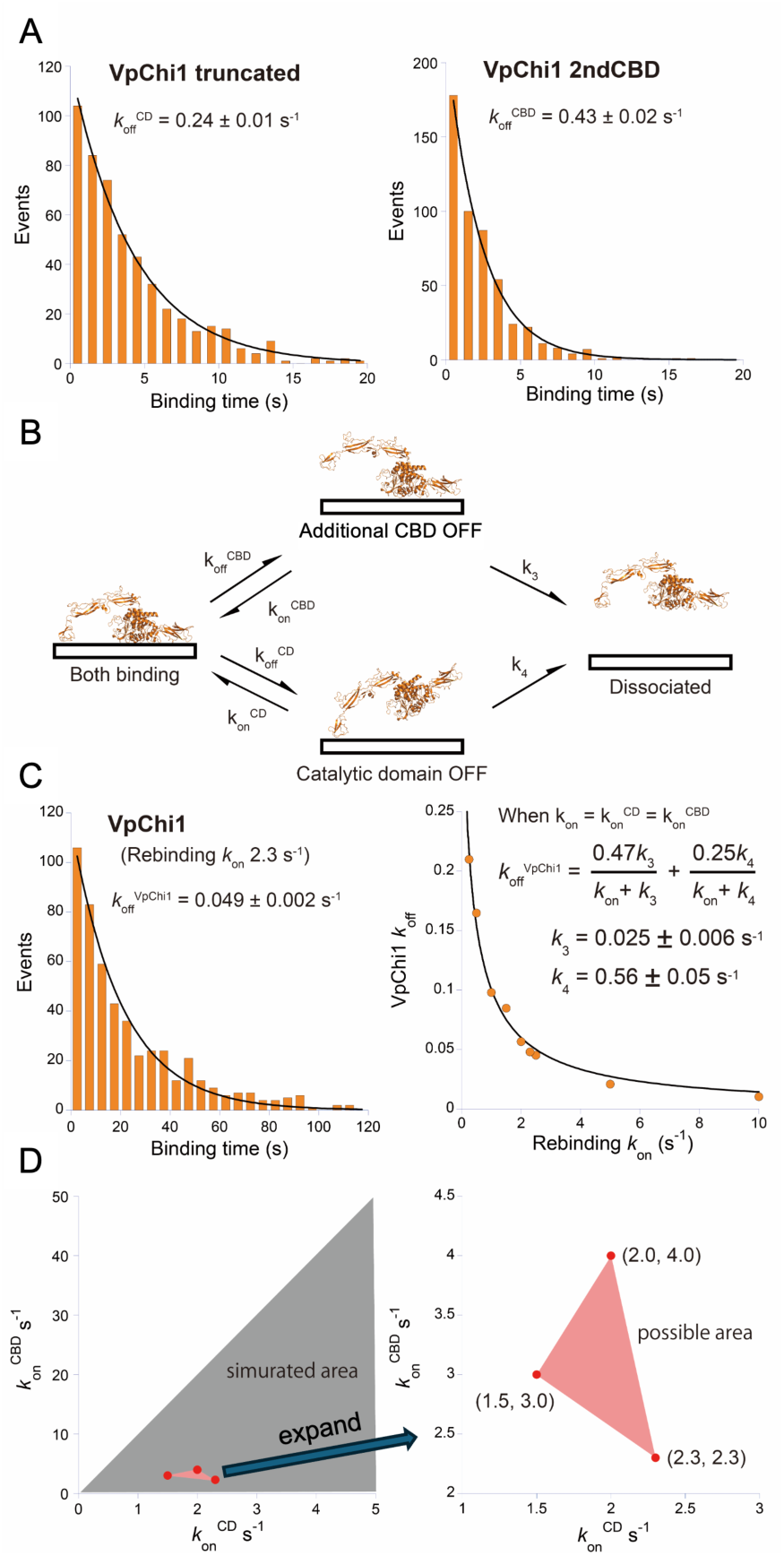
Binding-time simulation of VpChi1 on the hydrophobic surface. (A) Simulated binding-time distributions for VpChi1-truncated and the 2ndCBD on the hydrophobic surface, generated using the experimentally determined *k*off values. (B) Sequential dissociation model for VpChi1. (C) Simulated binding-time distribution of VpChi1 using a rebinding *k*on of 2.3 s^-1^, and the relationship between the rebinding *k*on and the resulting *k*off, fitted using the equation derived from the sequential dissociation model (Supplementary Fig. S5). (D) Simulated combinations of *k*on^CD^ (0.25 to 5 s^-1^) and *k*on^CBD^ (0.025 to 50 s^-1^), with parameter pairs that successfully reproduced the experimentally observed *k*off of VpChi1.

To identify plausible combinations of *k* ^CD^ and *k* ^CBD^, we generated binding-time distributions over a wide parameter space (*k*_on_ ^CD^ = 0.25–5 s⁻¹; *k* ^CBD^ = 0.025–50 s^-1^) (Fig. S4). The parameter sets that lay within the region enclosed by the three plots and best recapitulated the experimental behaviour were (*k* ^CD^, *k* ^CBD^) = (1.5, 3.0), (2.0, 4.0) and (2.3, 2.3) s^-1^ (Fig. 4D).

### Analysis of moving velocity and moving distance

The movement velocities and moving distances of VpChi1 A352C–Cy3 and VpChi1-truncated A352C–Cy3 were analysed using HS-AFM (Fig. S5). Both enzymes displayed smooth, processive motion along the surface of a single chitin crystal, and no additional binding of the extra domains in full-length VpChi1 was observed (Supplementary Movies 4 and 5). The velocity of VpChi1 was 51.5 nm s^-1^, whereas VpChi1-truncated moved at 47.9 nm s^-1^. The mean durations of individual movement events were 1.1 s for VpChi1 and 1.0 s for the truncated enzyme. These measurements indicate that VpChi1 moved approximately 38.4 nm along the crystal, while VpChi1-truncated moved 30.3 nm.

### Chitin degradation bacteria screening

Chitin-utilising bacteria were screened from seawater and marine sediments collected at a range of depths (100–4819 m) around northern and southern Japan. Marine bacteria that degraded chitin and produced pits on chitin plates were dominated by *Pseudoalteromonas* spp. and *Vibrio* spp., which were detected across nearly all sampling depths (Table 1). *Shewanella* sp., the third most frequently isolated genus, is a well-known degrader of chitin [23] as well as other polysaccharides [24]. Several minor taxa that formed pits on chitin plates have also been previously reported to exhibit chitin-utilisation capabilities. For example, *Thalassiosira pseudonana*, a member of the *Thalassospira* group, possesses 24 chitinase genes in its genome [25]. In the coastal Pacific Ocean, the majority of chitinase genes recovered from metagenomes have been attributed to the *Roseobacter* clade [26]. Genes involved in chitin utilisation have also been reported in eight strains of *Labrenzia alba* (formerly *Stappia alba*) [27]. As an exception, we found no published reports describing chitin-degrading activity in *Psychrobacter marincola*.

**Table 1.**
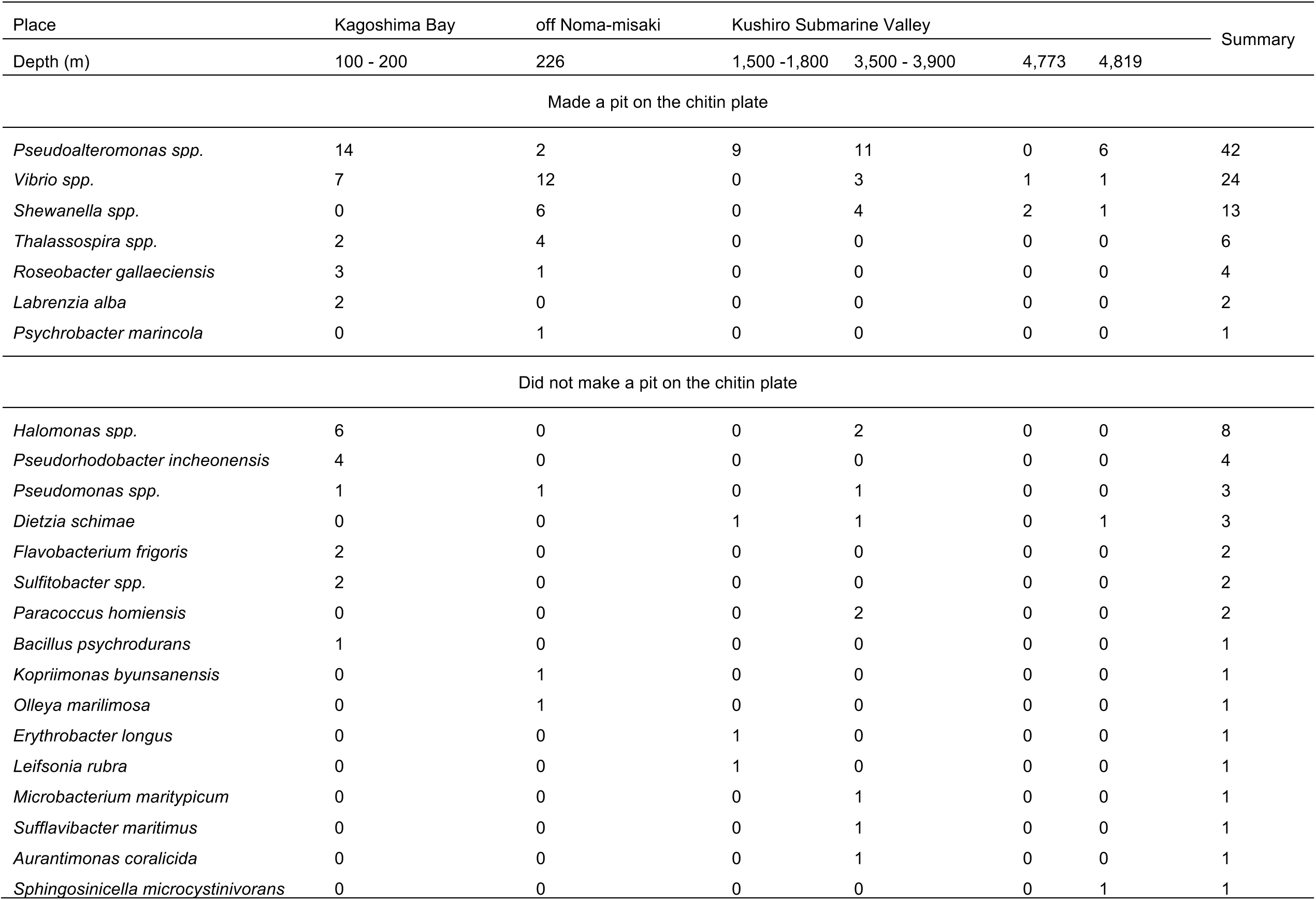
Identified bacteria from sea on chitin plate

Sixteen additional minor bacterial species that did not form visible pits on chitin plates nevertheless grew on the medium and were identified. Among these, *Hallomonas* sp. [28], *Pseudomonas* sp. [29], *Flavobacterium* sp. [30], *Paracoccus* sp. [31], marine *Bacillus* sp. [32], and *Psychrobacter* sp. [33] have been reported either to encode chitinases or to display chitin-degrading activity. Moreover, two chitosanases have been purified and characterised from *Microbacterium* sp. [34]. The remaining 13 species have not been reported to degrade chitin; however, they are common marine taxa and were likely present in the samples as background community members.

### Chitinase classification

The sequence diversity of GH18 chitinases from marine and non-marine bacteria was examined through phylogenetic analysis. Amino acid sequences of 270 chitinases similar to SmChiA or VpChi1 were retrieved using the NCBI BLASTp suite and aligned with the Constraint-based Multiple Alignment Tool (COBALT) after removal of predicted signal peptides identified with SignalP 5.0. A phylogenetic tree was constructed using SmChiB as an outgroup (Fig. 5). Notably, chitinases derived exclusively from marine bacteria formed a large, well-supported clade. This major marine clade was dominated by *Pseudoalteromonas* spp. and *Vibrio* spp., but also included *Enterovibrio*, *Grimontia*, *Aliivibrio fischeri*, and five *Photobacterium* spp., collectively designated as marine bacteria group 2. Two additional marine clades (groups 1 and 3) comprised seven *Photobacterium* spp., three *Vibrio* spp., *Endozoicomonas acroporae*, two *Aliivibrio* spp., two *Gallaecimonas* spp., and five *Microbulbifer* spp., respectively.

**Fig. 5.**
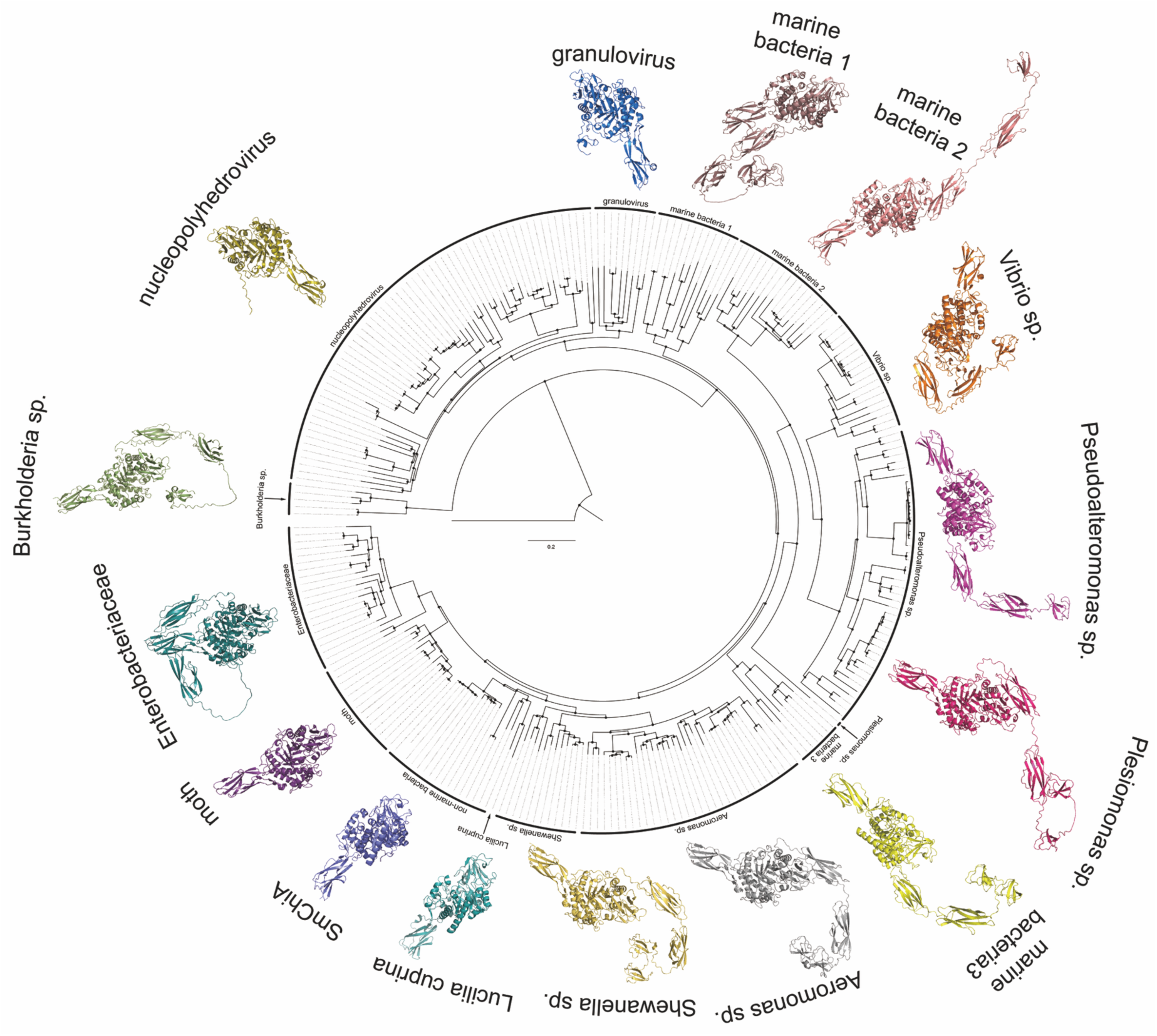
Guide tree and predicted structures of GH18 chitinases. A total of 270 GH18 chitinase sequences obtained from the NCBI BLASTp suite using SmChiA or VpChi1 as queries were aligned using the Constraint-based Multiple Alignment Tool (COBALT) (https://www.ncbi.nlm.nih.gov/tools/cobalt/). N-terminal signal peptides were removed prior to alignment based on predictions from SignalP 5.0 (https://services.healthtech.dtu.dk/service.php?SignalP-5.0). SmChiB (CAA85292.1) was included as an outgroup. Overall structural models were predicted using ColabFold.

Chitinases from *Burkholderia* sp. and *Plesiomonas shigelloides* formed distinct, isolated branches.

SmChiA grouped with the non-marine bacterial clade, which also contained several genera associated with human pathogenicity, including *Providencia*, *Rahnella*, *Klebsiella*, *Enterobacter* and *Serratia*. This clade additionally contained chitinases from *Pseudomonas* spp. and *Sanguibacter* spp. Interestingly, a chitinase from the Australian sheep blowfly *Lucilia cuprina* (XP_023299420.1) clustered within the same branch as this non-marine bacterial group.

A clade of chitinases from *Shewanella* spp., which were also isolated in our chitin-degradation screening, was positioned phylogenetically closer to SmChiA than to VpChi1. Chitinases from *Aeromonas* spp. fell within the same branch as the *Shewanella* group. Both genera are commonly encountered in aquatic environments. Another distinct cluster comprised chitinases from *Enterobacteriaceae*, forming a neighbouring branch to the moth chitinase group. This *Enterobacteriaceae* cluster included chitinases from *Leclercia* sp., *Erwinia teleogrylli*, *Enterobacter* sp., *Escherichia coli*, *Citrobacter* sp., *Pedobacter himalayensis*, *Klebsiella aerogenes*, *Lelliottia* sp. and *Franconibacter* spp.

Chitinases from *Baculoviridae* were clearly separated from both bacterial and insect-derived chitinases. Those from nucleopolyhedroviruses and granuloviruses formed distinct clades independent of the bacterial GH18 lineages.

### Structural features of chitinases

One representative sequence from each phylogenetic group was selected, and the corresponding structures were predicted using ColabFold (Fig. 5). The domain architectures of the chitinases are summarised in Fig. S6. Chitinases within the same phylogenetic group exhibited highly similar domain organisations. All marine bacterial chitinases possessed one or two additional chitin-binding domains (classified in carbohydrate binding module family 5: CBM5) located at the C-terminus of the SmChiA-like GH18 catalytic domain. In most cases, two FN3 domains were inserted between the catalytic domain and the CBM5 modules, with the exception of chitinases belonging to marine bacteria group 1.

Chitinases from the *Aeromonas* and *Enterobacteriaceae* groups also contained additional FN3 and CBM5 domains, resembling the architectures observed in marine bacterial chitinases. Although *Aeromonas* spp. were not isolated from marine samples in this study, they commonly inhabit freshwater and brackish environments. The remaining taxa occupy terrestrial habitats, where their chitinases are likely to operate under low-moisture conditions or on cellular surfaces rather than in fully aquatic systems.

## Discussion

VpChi1-truncated shares a highly similar structural architecture with SmChiA. Its *k*cat and Km values towards crystalline chitin (4.1 s^-1^ and 0.50 mg mL^-1^; Fig. 2) were both higher than those reported for SmChiA (*k*cat 3.1 s^-1^, Km 0.32 mg mL^-1^) [35]. The higher Km of VpChi1-truncated can be attributed to differences in the aromatic residues forming the flat binding surface. VpChi1 contains Tyr31, Trp70, Trp231 and Tyr245 (Fig. 1), whereas SmChiA possesses Trp69, Trp33, Trp245 and Phe232, resulting in a higher proportion of tryptophan residues. Because tryptophan side chains display ∼1.2-fold stronger binding free energy to hydrophobic surfaces than tyrosine [36], SmChiA exhibits tighter binding to crystalline chitin compared with VpChi1-truncated.

Full-length VpChi1 showed a higher *k*cat (5.4 s^-1^) and a ninefold lower Km (0.055 mg mL^-1^) relative to VpChi1-truncated, demonstrating that the additional CBD markedly enhances affinity for crystalline chitin. The catalytic efficiency (*k*cat/Km) of VpChi1 (98 s^-1^ mg^-1^ mL) was 12-fold higher than that of the truncated enzyme (8.2 s^-1^ mg^-1^ mL). However, at chitin concentrations above 1.5 mg mL^-1^, the activities of the two enzymes became similar. The most plausible explanation is that the additional CBD binds to neighbouring crystals, leading to substrate inhibition. Under these conditions, the enzyme may not be fully inactive but instead produces chitobiose more slowly. Although a modified kinetic model provided improved fitting (Fig. S2), the large fitting errors arising from a high degree of freedom in *k*cat2 limit its interpretive value.

To understand how the additional CBD enhances affinity, it is necessary to examine conditions where the effective chitin concentration is low. In single-molecule HS-AFM and fluorescence microscopy, we observed individual enzyme molecules interacting with single (or bundled) crystals, minimising crowding effects. Surprisingly, the apparent *k*on values of VpChi1 (5.6 × 10^8^ μm^-1^ s^-1^ M^-1^) and VpChi1-truncated (4.5 × 10^8^ μm^-1^ s^-1^ M^-1^) were very similar. The most prominent difference lay in surface selectivity. VpChi1 possesses a flat hydrophobic-binding surface similar to SmChiA [13], and the slow-dissociation state corresponds to binding on this hydrophobic face. Apparent *k*on and *k*off values for hydrophobic and hydrophilic binding states are summarised in Table 2.

**Table 2.**
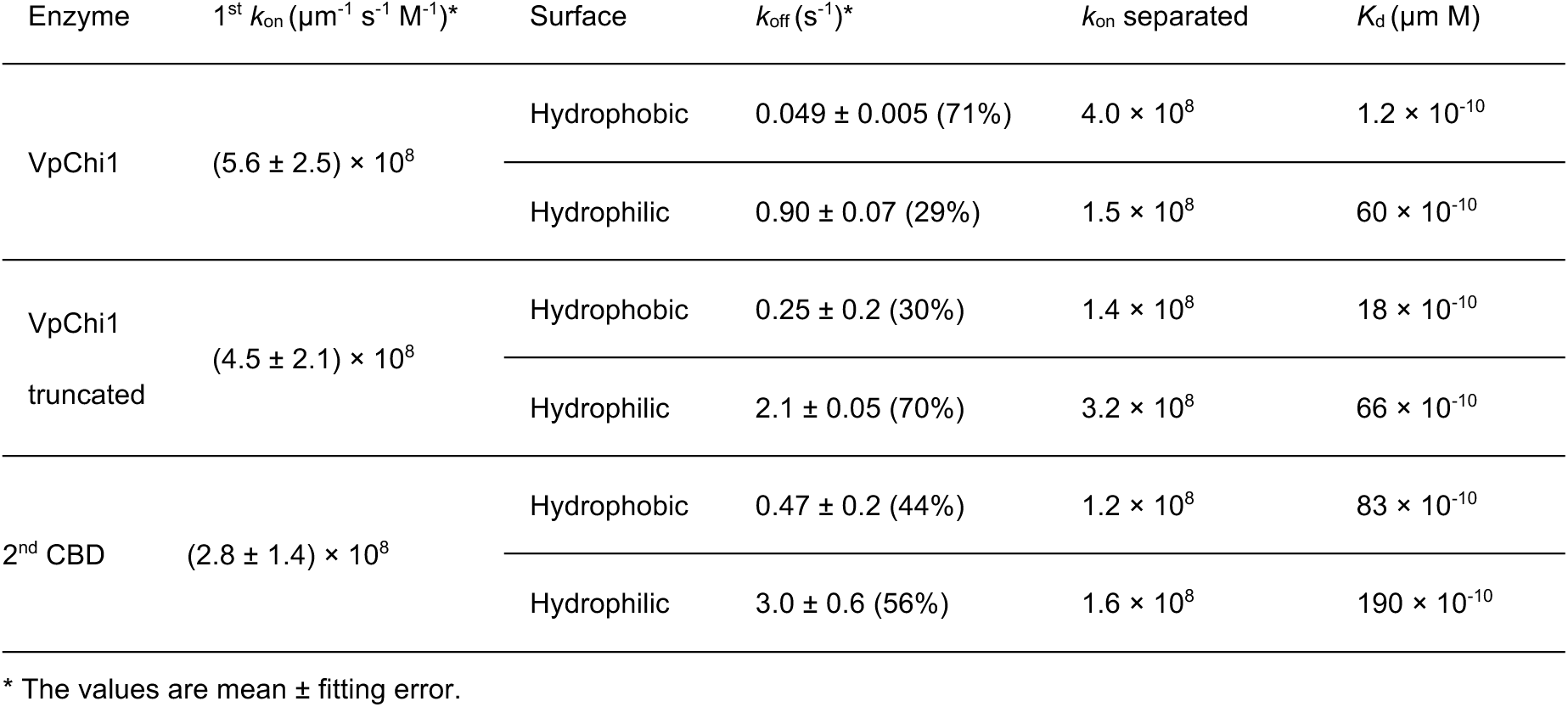
Summary of binding parameters of VpChi1 and truncated mutant

Deconvolution of the binding populations revealed that 71% of VpChi1 molecules resided on hydrophobic surfaces, 2.4-fold higher than VpChi1-truncated. VpChi1 exhibited a hydrophobic-surface *k*on of 4.0 × 10^8^ μm^-1^ s^-1^ M^-1^, compared with 1.4 × 10^8^ μm^-1^ s^-1^ M^-1^ for VpChi1-truncated. Furthermore, the *k*off from hydrophobic surfaces for VpChi1 (0.049 s⁻¹) was 5.1-fold lower than that of VpChi1-truncated (0.25 s^-1^). Consequently, the resulting Kd values were 1.2 × 10^-10^ μm M for VpChi1 and 18 × 10^-10^ μm M for VpChi1-truncated, a 15-fold difference, mirroring the magnitude of the *k*cat/Km relationship. By contrast, affinities for hydrophilic surfaces were similar (60 × 10^-10^ μm M vs. 66 × 10^-10^ μm M), indicating that the additional CBD does not enhance binding to hydrophilic faces.

To further characterise the role of the CBD, we examined the isolated 2ndCBD. Its apparent *k*on was 62% of VpChi1-truncated, yet its hydrophobic-surface *k*on (1.2 × 10^8^ μm^-1^ s^-1^ M^-1^) was similar to the truncated enzyme. Its *k*off from hydrophobic surfaces (0.47 s^-1^) was 1.9-fold faster than VpChi1-truncated. Given that VpChi1 remains bound to hydrophobic surfaces for ∼20 s, 5 to 9.5 times longer than either VpChi1-truncated (4.0 s) or 2ndCBD (2.1 s), the two domains appear to act synergistically. Simulated binding-time distributions reproduced the VpChi1 profile when a sequential dissociation model was applied, and the closest match to experimental *k*off was obtained with a rebinding rate constant of 2.3 s^-1^. These findings indicate that cooperativity arises because one domain facilitates rebinding of the other. Introducing a second binding domain to VpChi1-truncated can reduce total *k*off fivefold through this rebinding assistance. Moreover, because *k*3 (CD dissociation after CBD dissociation) was 18-fold smaller than *k*4 (CBD dissociation after CD dissociation), VpChi1 dissociation primarily occurs through CD release following CBD dissociation. Parameter combinations that reproduced VpChi1-like *k*off values required *k*on^CBD^ to be approximately twice *k*on^CD^, indicating that balanced adsorption–desorption kinetics between domains underpin domain cooperativity.

Screening of marine bacteria identified *Pseudoalteromonas*, *Vibrio* and *Shewanella* spp. as the dominant chitin degraders, capable of using chitin as the sole carbon source. Because chitin polymers are too large for direct uptake, growth required extracellular degradation—consistent with the GH18 chitinase clades recovered in our phylogeny (Fig. 5). Although *Halomonas* sp. has reported chitinase activity [25], no GH18 sequence was detected in BLAST searches, and it was therefore excluded from the phylogeny. In GH18 chitinases resembling SmChiA, the additional CBDs were consistently C-terminal CBM5 modules linked by FN3 domains, with the exception of marine group 1, which partly used peptide linkers. A SmChiB-like architecture (GH18 CD + C-terminal CBM5) was not observed among marine chitinases; instead, they resembled a fusion of SmChiA and SmChiB. In HS-AFM, SmChiA and SmChiB move in opposite directions on chitin [10], whereas VpChi1 and VpChi1-truncated moved in the same direction, indicating that the additional CBD does not influence directional movement and may instead remain flexible or swing freely during catalysis.

The primary role of the additional CBD is therefore to prolong CD adsorption to hydrophobic crystal faces. This architecture may be highly conserved among aquatic bacteria because, in water, enzymes readily diffuse away upon dissociation. For chemotactic taxa such as *Vibrio* spp., maintaining prolonged contact with chitin enables continuous production of chitobiose, thereby providing a persistent chemical signal that guides cells to the particle (Fig. 6). In contrast, terrestrial environments have limited water and high effective substrate concentrations; under such conditions, excessive adsorption can impede turnover more than diffusion loss, favouring chitinases without additional CBDs, such as those of Serratia marcescens. This represents a striking example of how enzyme structural features reflect microbial ecological niches.

**Fig. 6.**
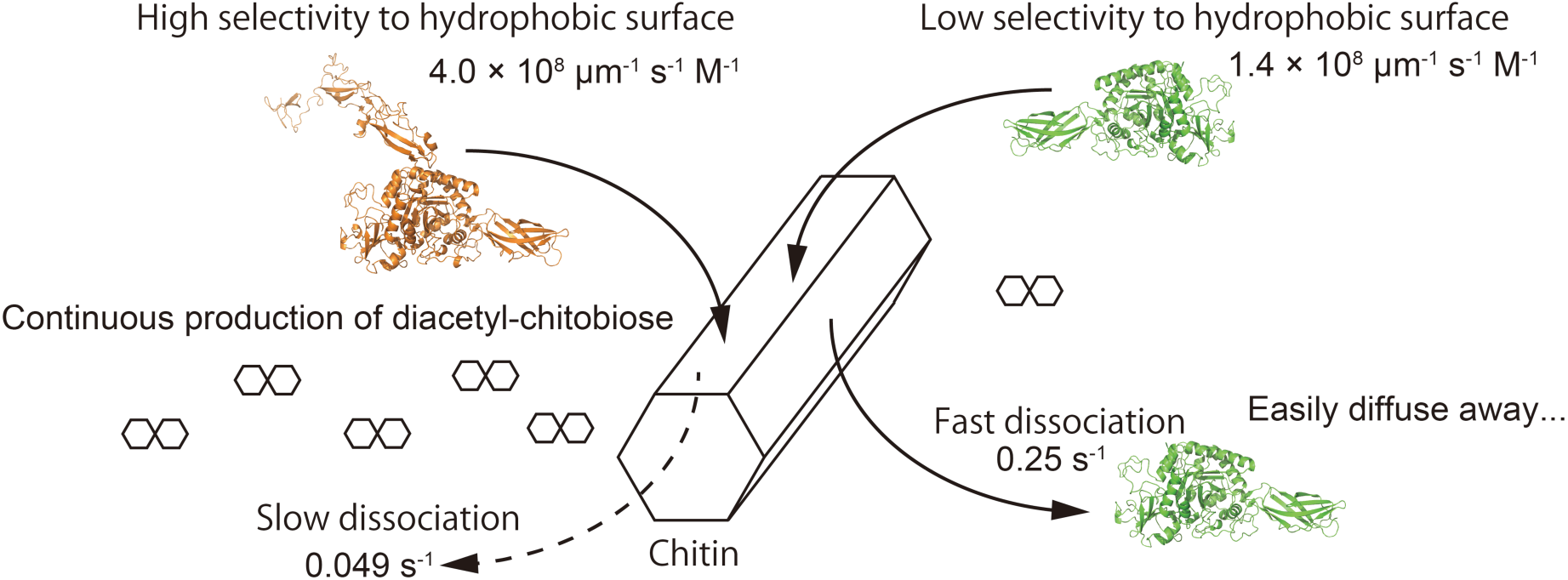
Biological significance of double CBD architecture in marine bacteria chitinases. The additional CBD enhances binding specificity to the degradable hydrophobic surface of crystalline chitin and decreases the *k*off, thereby enabling continuous production of diacetylchitobiose. This sustained production provides a persistent chemical signal that allows marine bacteria to localise to chitin-rich sites.

## Study funding

This study was financially supported by MEXT Leading Initiative for Excellent Young Researchers Grant Number 201990171 (to A. N.), JSPS KAKENHI Grant Number JP19H03094 (to A. N.), and Technology of Japan and JST FOREST Program (Grant Number JPMJFR210C to A. N.).

## Supporting information

MovieS1_VpChi1_x10play

MovieS2_VpChi1-truncated_x10play

MovieS3_VpChi1-2ndCBD_x10play

MovieS4_VpChi1_x3play

MovieS5_VpChi1-truncated_x3play

## Abbreviations

catalytic domain CD; chitin-binding domain CBD; glycoside hydrolase families GH; diacetylchitobiose GlcNAc₂; high-speed atomic force microscopy HS-AFM; fibronectin type III-like FN3: carbohydrate binding module family 5 CBM5;

## Notes

### Competing Interest Statement

The authors have declared no competing interest.

