## Supplemental data for "A dual-domain chitinase mechanism enables marine bacteria to sense and localize sparse crystalline chitin in the ocean"

Contents

Table S1. Data collection and refinement statistics of VpChi1-truncated
Table S2. Detailed data of chitin-degrading bacteria screening.
(Attached as a different file in Excel format)
Fig. S1 Activity plots against crystalline chitin concentration in artificial sea water at pH7.5.
Fig. S2 Curve fitting with the model including a diacetyl-chitobiose production from the double binding state.
Fig. S3 Dissociation model and equation between $k_{off}$ and rebinding $k_{on}$.
Fig. S4 Binding time distribution simulations with different combinations of $k_{on}$ values
Fig. S5 Distributions of moving velocities and moving length of VpChi1-A352C-Cy3 and the truncated mutant analyzed by HS-AFM.
Fig. S6 Domain compositions and estimated structures of the chitinases.
Fig. S7 Python script for binding time simulation of single domain.
Fig. S8 Python script for binding time simulation of double domain.

Legends for Supplementary movies

### Table S1. Data collection and refinement statistics of VoChi1-truncated.

| | VpChi1-truncated |
|---|---|
| PDB ID | 8HRF |
| Wavelength | 1.54 |
| Resolution range | 28.7 - 2.5 (2.589 - 2.5) |
| Space group | P 1 21 1 |
| Unit cell | 48.8042 159.996 75.4886 90 92.6828 90 |
| Total reflections | 287428 (29979) |
| Unique reflections | 39898 (3994) |
| Multiplicity | 7.2 (7.5) |
| Completeness (%) | 98.75 (97.42) |
| Mean I/sigma(I) | 13.30 (5.63) |
| Wilson B-factor | 14.17 |
| R-int | 0.149 (0.330) |
| Reflections used in refinement | 39441 (3894) |
| Reflections used for R-free | 2021 (203) |
| R-work | 0.1659 (0.1776) |
| R-free | 0.2353 (0.2797) |
| Number of non-hydrogen atoms | 9435 |
| macromolecules | 8710 |
| ligands | 17 |
| solvent | 714 |
| Protein residues | 1138 |
| RMS(bonds) | 0.008 |
| RMS(angles) | 0.90 |
| Ramachandran favored (%) | 95.06 |
| Ramachandran allowed (%) | 4.76 |
| Ramachandran outliers (%) | 0.18 |
| Rotamer outliers (%) | 3.44 |
| Clash score | 12.83 |
| Average B-factor | 17.76 |
| macromolecules | 17.58 |
| ligands | 19.78 |
| solvent | 19.94 |

Statistics for the highest-resolution shell are shown in parentheses.

### Table S2. Sampling conditions of chitin degrading bacteria.

| Cruise no. | Dive no. | Place | Depth | Latitude | Longitude | Related species (16S rRNA sequence identity, %) of Isolated strains |
|---|---|---|---|---|---|---|
| NT07-09 | 682 | off Noma-misaki | 226 | 31º20.725N | 129º59.289E | *Vibrio pomeroyi* (100), *Shewanella japonica* (98–99), *Vibrio splendidus* (99–100), *Thalassospira lucentensis* (99), *Vibrio cyclitrophicus* (98), *Pseudoalteromonas piscicida* (100), *Psychrobacter marincola* (99), |
| NT07-09 | 683 | Kagoshima Bay | 200 | 31º39.506N | 130º46.485E | *Pseudoalteromonas piscicida* (99–100), *Pseudoalteromonas sp* (100), *Roseobacter gallaeciensis* (95), *Seudoalteromonas piscicida* (100), *Labrenzia alba* (98), *Vibrio fischeri* (99), *Vibrio splendidus* (99), |
| NT07-09 | 686 | Kagoshima Bay | 102 | 31º39.742N | 130º48.052E | *Vibrio splendidus* (98), *Pseudoalteromonas elyakovii* (99), *Vibrio rumoiensis* (99), *Vibrio tasmaniensis* (98), *Pseudoalteromonas arctica* (100), |
| YK07-14 | 1029 | Kushiro Submarine Valley | 4,773 | 41°15.331N | 144°38.549E | *Pseudoalteromonas elyakovii* (99), *Pseudoalteromonas sp* (100),*Pseudoalteromonas haloplanktis* (100), *Pseudoalteromonas mariniglutinosa* (100), *Pseudoalteromonas whanghaensis* (98), *Shewanella oneidensis* (95), *Vibrio splendidus* (100), *Vibrio campbellii* (99), |
| YK07-14 | 1030 | Kushiro Submarine Valley | 4,819 | 41°15.293N | 144°38.578E | *Shewanella baltica* (100), *Vibrio campbellii* (100), |
| YK07-14 | 1031 | Kushiro Submarine Valley | 3,510 | 41°16.094N | 144°34.672E | *Pseudoalteromonas whanghaensis* (99), *Vibrio splendidus* (100), |
| YK07-14 | 1032 | Kushiro Submarine Valley | 3,556 | 41°15.957N | 144°34.562E | *Pseudoalteromonas whanghaensis* (99), |
| YK07-14 | 1033 | Kushiro Submarine Valley | 1,515 | 42°29.201N | 144°39.157E | *Pseudoalteromonas whanghaensis* (99–100), *Shewanella putrefaciens* (99), |
| YK07-14 | 1034 | Kushiro Submarine Valley | 1,521 | 42°28.899N | 144°39.398E | *Pseudoalteromonas ganghwensis* (96–100), *Pseudoalteromonas whanghaensis* (98), |
| YK07-14 | 1035 | Kushiro Submarine Valley | 3,750 | 42°03.671N | 145°09.653E | *Pseudoalteromonas ganghwensis* (99), *Pseudoalteromonas mariniglutinosa* (99), |

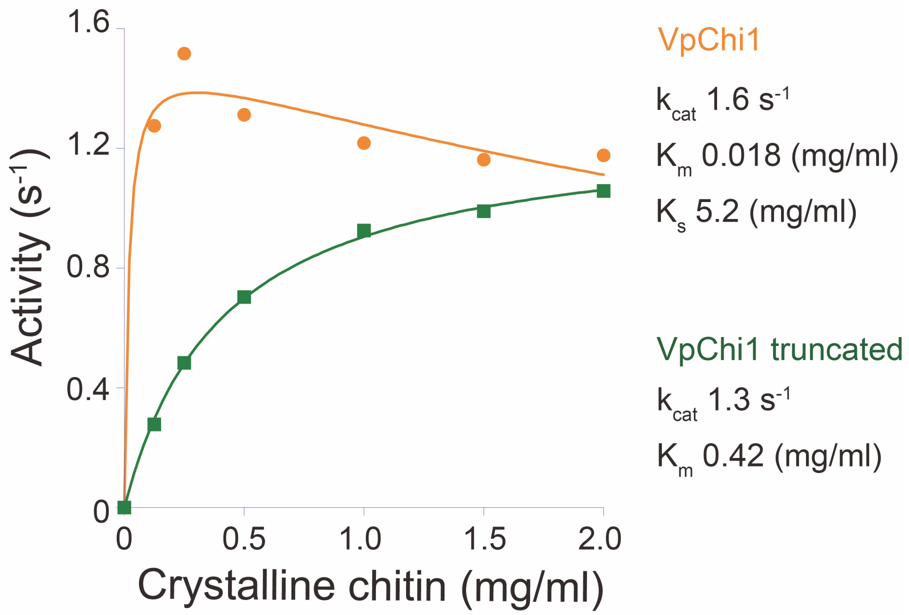


**Fig. S1 Activity plots against crystalline chitin concentration in artificial sea water at pH7.5.**

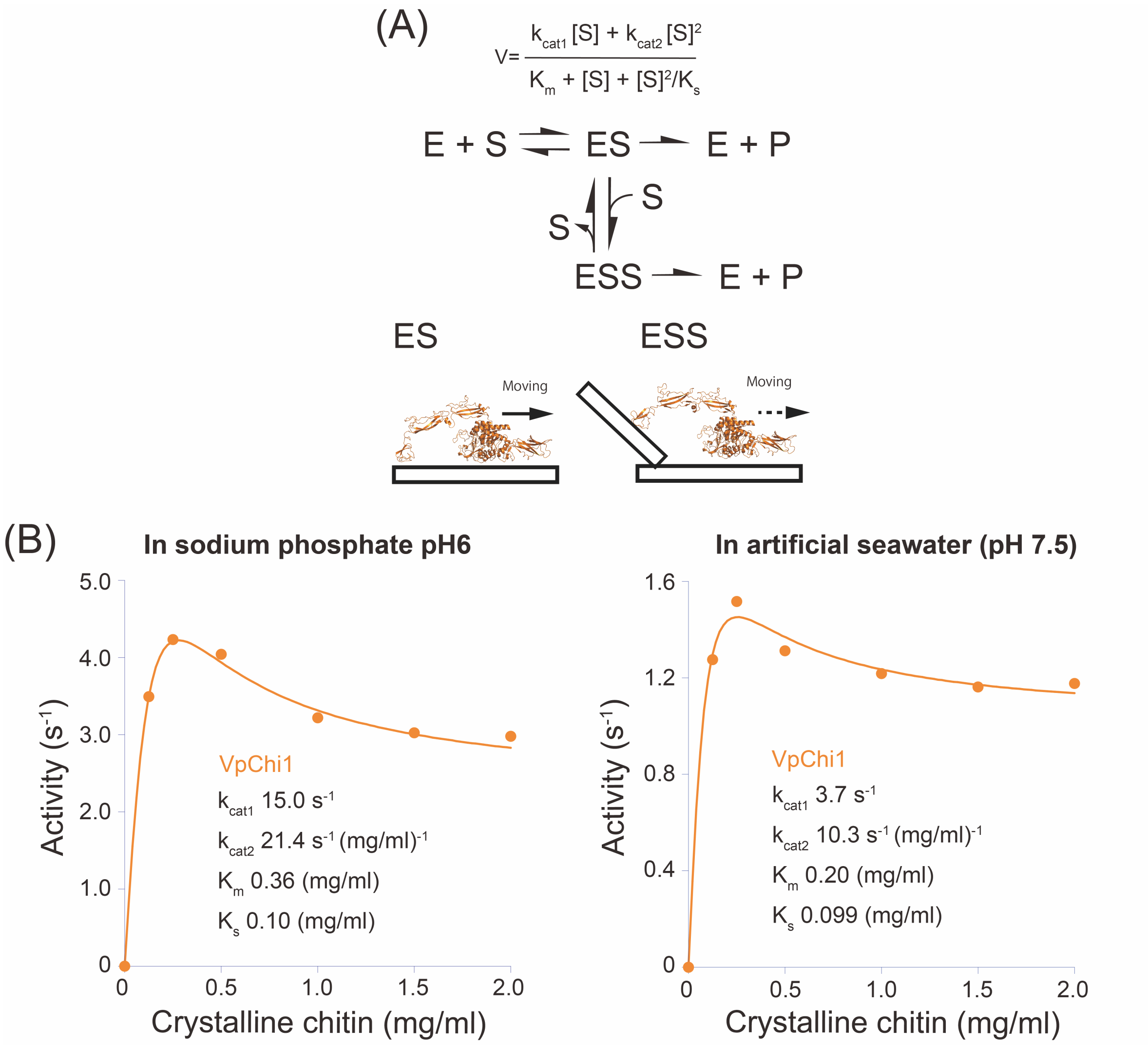


**Fig. S2 Curve fitting with the model including a diacetyl-chitobiose production from the double binding state.**

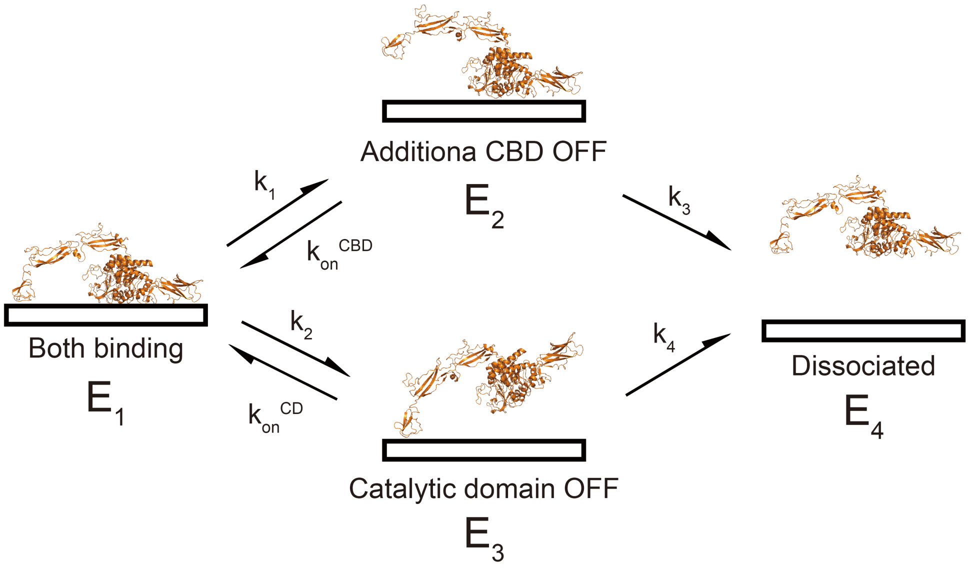


What we want to know is $k_{off}$ of VpChi1

$$\frac{dE_4}{dt} = k_{off}E_1 = k_3E_2 + k_4E_3$$

Here,

$$\frac{dE_2}{dt} = k_1E_1 - (k_{on}^{CBD} + k_3)E_2 \quad \frac{dE_3}{dt} = k_2E_1 - (k_{on}^{CD} + k_4)E_3$$

When we observe enough numbers of molecules, it looks like an equilibrium state

$$\frac{dE_2}{dt} = 0 \quad \frac{dE_3}{dt} = 0 \qquad E_2 = \frac{k_1E_1}{k_{on}^{CBD} + k_3} \quad E_3 = \frac{k_2E_1}{k_{on}^{CD} + k_4}$$

Therefore,

$$k_{off} = \frac{k_1k_3}{k_{on}^{CBD} + k_3} + \frac{k_2k_4}{k_{on}^{CD} + k_4}$$

**Fig. S3 Dissociation model and equation between $k_{off}$ and rebinding $k_{on}$.**

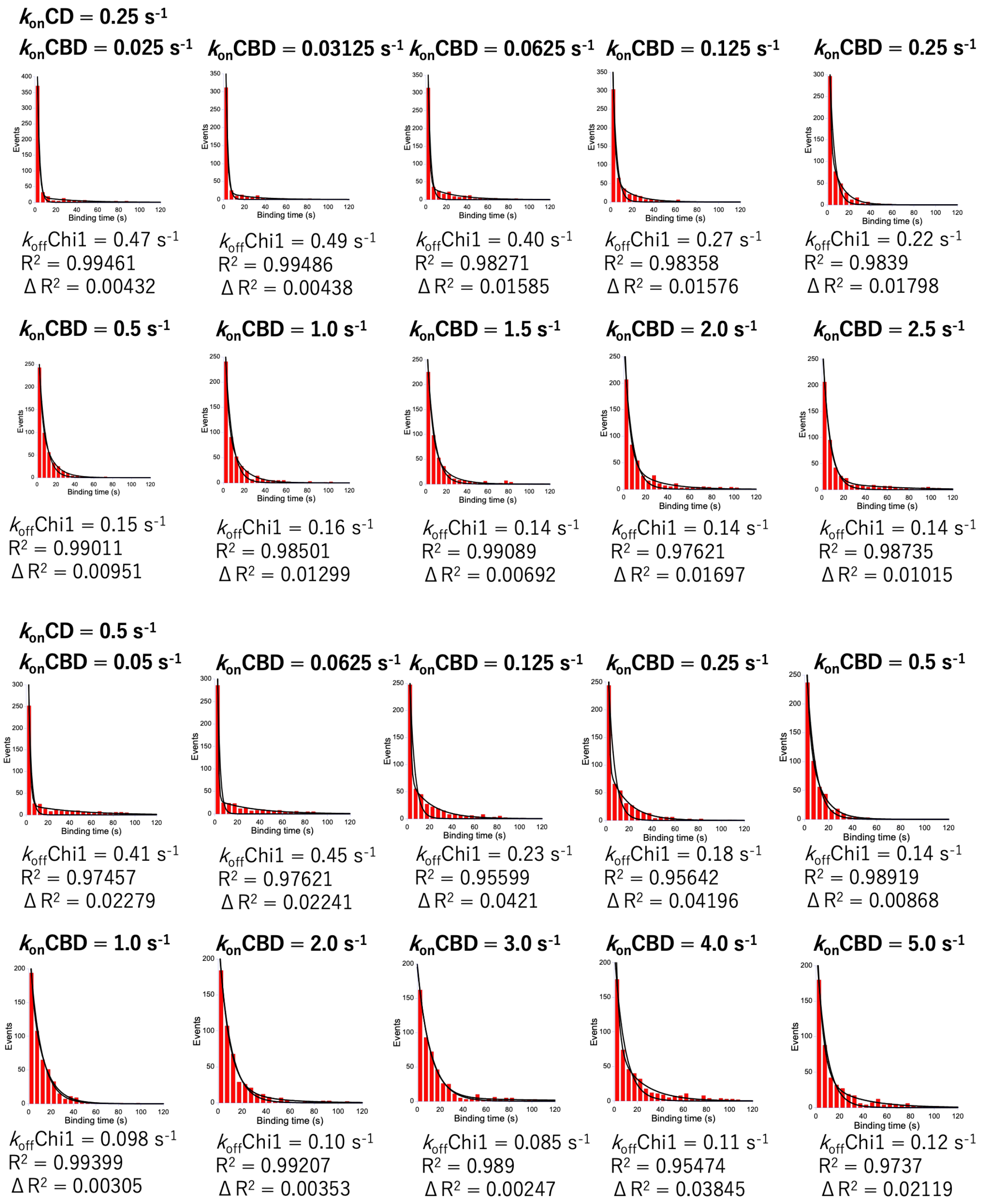

$k_{on}$CD = 0.25 s$^{-1}$
$k_{on}$CBD = 0.025 s$^{-1}$
$k_{on}$CBD = 0.03125 s$^{-1}$
$k_{on}$CBD = 0.0625 s$^{-1}$
$k_{on}$CBD = 0.125 s$^{-1}$
$k_{on}$CBD = 0.25 s$^{-1}$
Events
Binding time (s)
$k_{off}$Chi1 = 0.47 s$^{-1}$
$R^2$ = 0.99461
$\Delta R^2$ = 0.00432
$k_{off}$Chi1 = 0.49 s$^{-1}$
$R^2$ = 0.99486
$\Delta R^2$ = 0.00438
$k_{off}$Chi1 = 0.40 s$^{-1}$
$R^2$ = 0.98271
$\Delta R^2$ = 0.01585
$k_{off}$Chi1 = 0.27 s$^{-1}$
$R^2$ = 0.98358
$\Delta R^2$ = 0.01576
$k_{off}$Chi1 = 0.22 s$^{-1}$
$R^2$ = 0.9839
$\Delta R^2$ = 0.01798
$k_{on}$CBD = 0.5 s$^{-1}$
$k_{on}$CBD = 1.0 s$^{-1}$
$k_{on}$CBD = 1.5 s$^{-1}$
$k_{on}$CBD = 2.0 s$^{-1}$
$k_{on}$CBD = 2.5 s$^{-1}$
$k_{off}$Chi1 = 0.15 s$^{-1}$
$R^2$ = 0.99011
$\Delta R^2$ = 0.00951
$k_{off}$Chi1 = 0.16 s$^{-1}$
$R^2$ = 0.98501
$\Delta R^2$ = 0.01299
$k_{off}$Chi1 = 0.14 s$^{-1}$
$R^2$ = 0.99089
$\Delta R^2$ = 0.00692
$k_{off}$Chi1 = 0.14 s$^{-1}$
$R^2$ = 0.97621
$\Delta R^2$ = 0.01697
$k_{off}$Chi1 = 0.14 s$^{-1}$
$R^2$ = 0.98735
$\Delta R^2$ = 0.01015
$k_{on}$CD = 0.5 s$^{-1}$
$k_{on}$CBD = 0.05 s$^{-1}$
$k_{on}$CBD = 0.0625 s$^{-1}$
$k_{on}$CBD = 0.125 s$^{-1}$
$k_{on}$CBD = 0.25 s$^{-1}$
$k_{on}$CBD = 0.5 s$^{-1}$
$k_{off}$Chi1 = 0.41 s$^{-1}$
$R^2$ = 0.97457
$\Delta R^2$ = 0.02279
$k_{off}$Chi1 = 0.45 s$^{-1}$
$R^2$ = 0.97621
$\Delta R^2$ = 0.02241
$k_{off}$Chi1 = 0.23 s$^{-1}$
$R^2$ = 0.95599
$\Delta R^2$ = 0.0421
$k_{off}$Chi1 = 0.18 s$^{-1}$
$R^2$ = 0.95642
$\Delta R^2$ = 0.04196
$k_{off}$Chi1 = 0.14 s$^{-1}$
$R^2$ = 0.98919
$\Delta R^2$ = 0.00868
$k_{on}$CBD = 1.0 s$^{-1}$
$k_{on}$CBD = 2.0 s$^{-1}$
$k_{on}$CBD = 3.0 s$^{-1}$
$k_{on}$CBD = 4.0 s$^{-1}$
$k_{on}$CBD = 5.0 s$^{-1}$
$k_{off}$Chi1 = 0.098 s$^{-1}$
$R^2$ = 0.99399
$\Delta R^2$ = 0.00305
$k_{off}$Chi1 = 0.10 s$^{-1}$
$R^2$ = 0.99207
$\Delta R^2$ = 0.00353
$k_{off}$Chi1 = 0.085 s$^{-1}$
$R^2$ = 0.989
$\Delta R^2$ = 0.00247
$k_{off}$Chi1 = 0.11 s$^{-1}$
$R^2$ = 0.95474
$\Delta R^2$ = 0.03845
$k_{off}$Chi1 = 0.12 s$^{-1}$
$R^2$ = 0.9737
$\Delta R^2$ = 0.02119

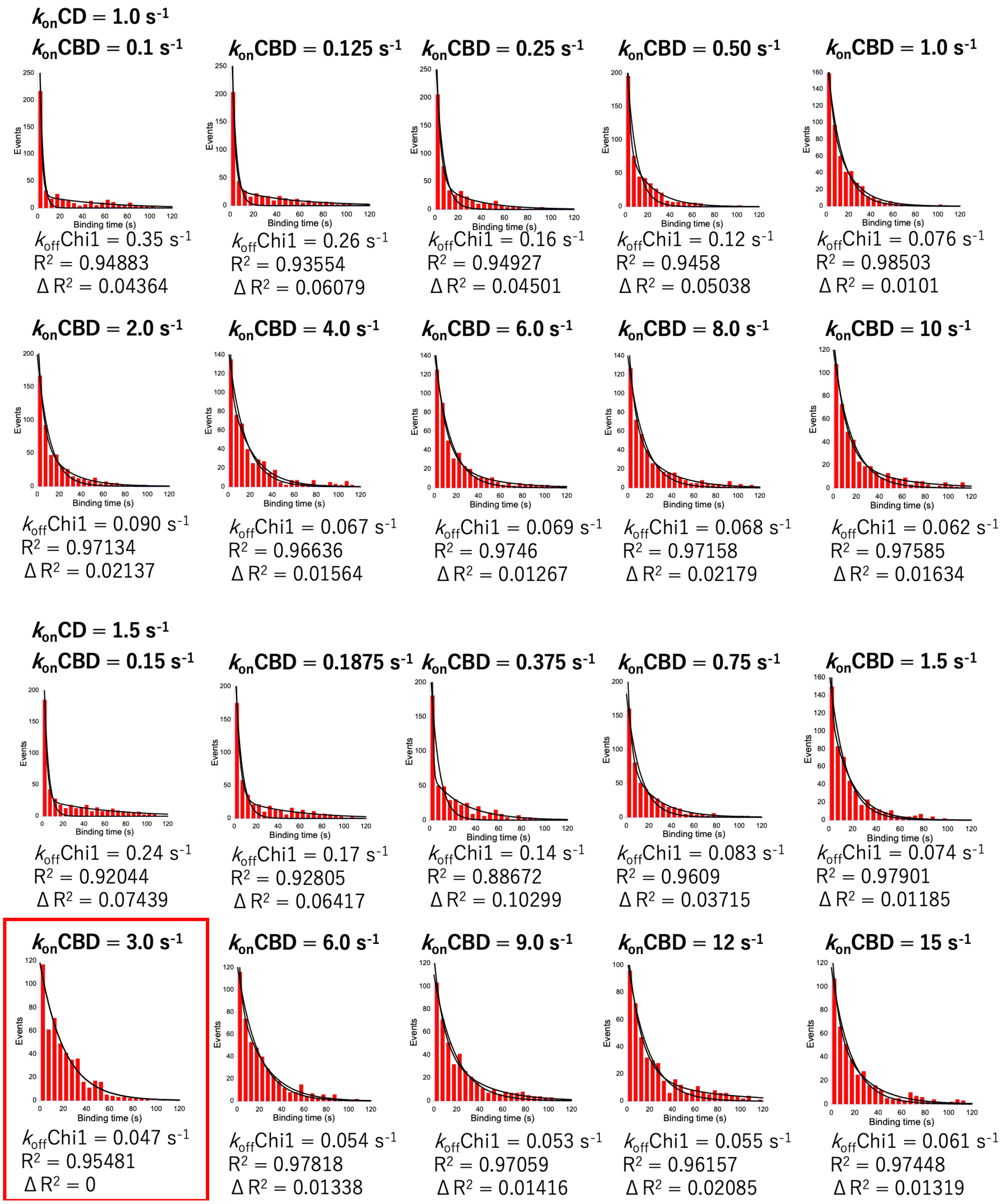
$k_{on}$CD = 1.0 $s^{-1}$
$k_{on}$CBD = 0.1 $s^{-1}$
$k_{off}$Chi1 = 0.35 $s^{-1}$
$R^2$ = 0.94883
Δ $R^2$ = 0.04364
$k_{on}$CBD = 0.125 $s^{-1}$
$k_{off}$Chi1 = 0.26 $s^{-1}$
$R^2$ = 0.93554
Δ $R^2$ = 0.06079
$k_{on}$CBD = 0.25 $s^{-1}$
$k_{off}$Chi1 = 0.16 $s^{-1}$
$R^2$ = 0.94927
Δ $R^2$ = 0.04501
$k_{on}$CBD = 0.50 $s^{-1}$
$k_{off}$Chi1 = 0.12 $s^{-1}$
$R^2$ = 0.9458
Δ $R^2$ = 0.05038
$k_{on}$CBD = 1.0 $s^{-1}$
$k_{off}$Chi1 = 0.076 $s^{-1}$
$R^2$ = 0.98503
Δ $R^2$ = 0.0101
$k_{on}$CBD = 2.0 $s^{-1}$
$k_{off}$Chi1 = 0.090 $s^{-1}$
$R^2$ = 0.97134
Δ $R^2$ = 0.02137
$k_{on}$CBD = 4.0 $s^{-1}$
$k_{off}$Chi1 = 0.067 $s^{-1}$
$R^2$ = 0.96636
Δ $R^2$ = 0.01564
$k_{on}$CBD = 6.0 $s^{-1}$
$k_{off}$Chi1 = 0.069 $s^{-1}$
$R^2$ = 0.9746
Δ $R^2$ = 0.01267
$k_{on}$CBD = 8.0 $s^{-1}$
$k_{off}$Chi1 = 0.068 $s^{-1}$
$R^2$ = 0.97158
Δ $R^2$ = 0.02179
$k_{on}$CBD = 10 $s^{-1}$
$k_{off}$Chi1 = 0.062 $s^{-1}$
$R^2$ = 0.97585
Δ $R^2$ = 0.01634
$k_{on}$CD = 1.5 $s^{-1}$
$k_{on}$CBD = 0.15 $s^{-1}$
$k_{off}$Chi1 = 0.24 $s^{-1}$
$R^2$ = 0.92044
Δ $R^2$ = 0.07439
$k_{on}$CBD = 0.1875 $s^{-1}$
$k_{off}$Chi1 = 0.17 $s^{-1}$
$R^2$ = 0.92805
Δ $R^2$ = 0.06417
$k_{on}$CBD = 0.375 $s^{-1}$
$k_{off}$Chi1 = 0.14 $s^{-1}$
$R^2$ = 0.88672
Δ $R^2$ = 0.10299
$k_{on}$CBD = 0.75 $s^{-1}$
$k_{off}$Chi1 = 0.083 $s^{-1}$
$R^2$ = 0.9609
Δ $R^2$ = 0.03715
$k_{on}$CBD = 1.5 $s^{-1}$
$k_{off}$Chi1 = 0.074 $s^{-1}$
$R^2$ = 0.97901
Δ $R^2$ = 0.01185
$k_{on}$CBD = 3.0 $s^{-1}$
$k_{off}$Chi1 = 0.047 $s^{-1}$
$R^2$ = 0.95481
Δ $R^2$ = 0
$k_{on}$CBD = 6.0 $s^{-1}$
$k_{off}$Chi1 = 0.054 $s^{-1}$
$R^2$ = 0.97818
Δ $R^2$ = 0.01338
$k_{on}$CBD = 9.0 $s^{-1}$
$k_{off}$Chi1 = 0.053 $s^{-1}$
$R^2$ = 0.97059
Δ $R^2$ = 0.01416
$k_{on}$CBD = 12 $s^{-1}$
$k_{off}$Chi1 = 0.055 $s^{-1}$
$R^2$ = 0.96157
Δ $R^2$ = 0.02085
$k_{on}$CBD = 15 $s^{-1}$
$k_{off}$Chi1 = 0.061 $s^{-1}$
$R^2$ = 0.97448
Δ $R^2$ = 0.01319
Events
Binding time (s)

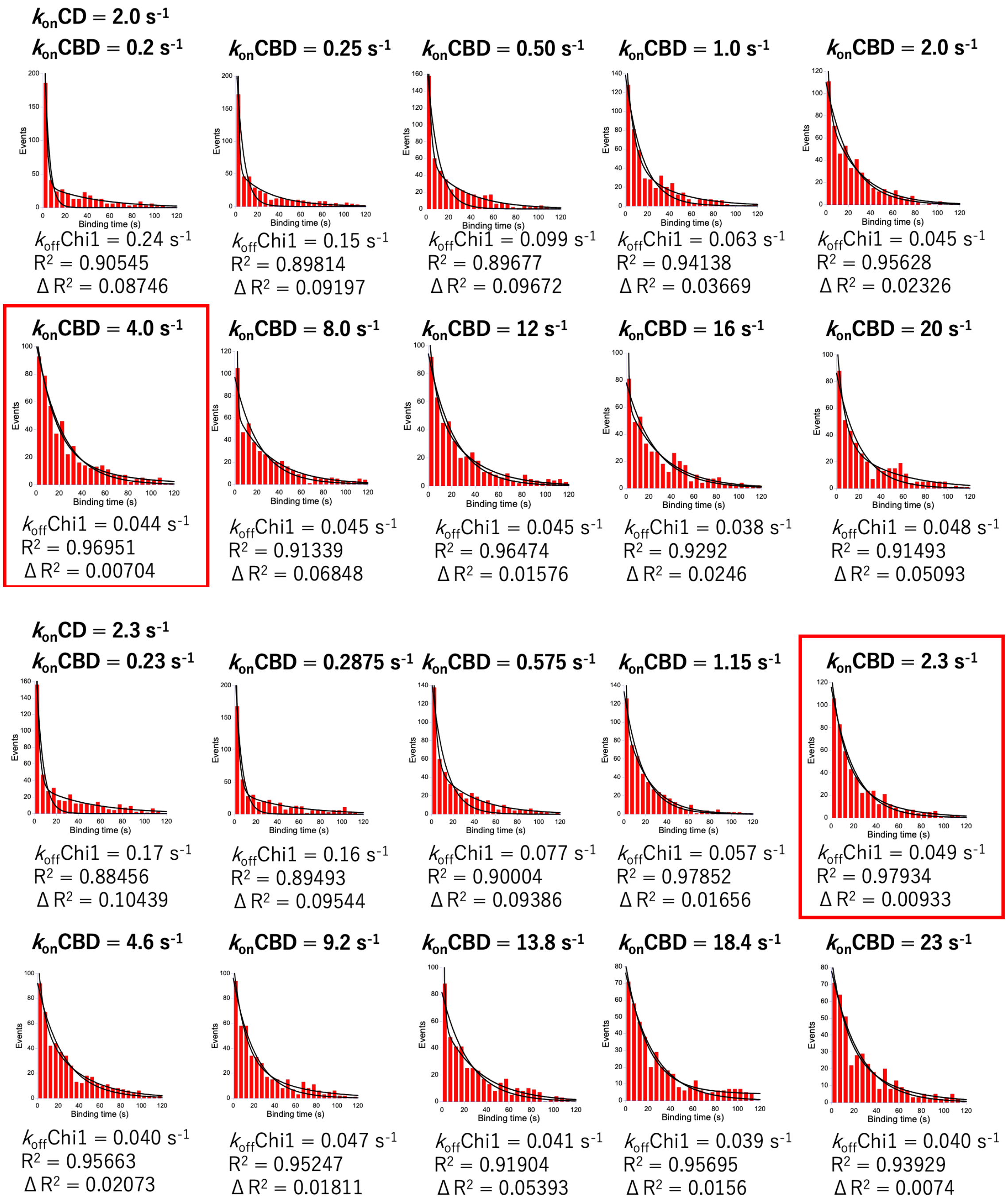

$k_{on}$CD = 2.0 $s^{-1}$
$k_{on}$CBD = 0.2 $s^{-1}$
$k_{off}$Chi1 = 0.24 $s^{-1}$
$R^2$ = 0.90545
Δ $R^2$ = 0.08746
$k_{on}$CBD = 0.25 $s^{-1}$
$k_{off}$Chi1 = 0.15 $s^{-1}$
$R^2$ = 0.89814
Δ $R^2$ = 0.09197
$k_{on}$CBD = 0.50 $s^{-1}$
$k_{off}$Chi1 = 0.099 $s^{-1}$
$R^2$ = 0.89677
Δ $R^2$ = 0.09672
$k_{on}$CBD = 1.0 $s^{-1}$
$k_{off}$Chi1 = 0.063 $s^{-1}$
$R^2$ = 0.94138
Δ $R^2$ = 0.03669
$k_{on}$CBD = 2.0 $s^{-1}$
$k_{off}$Chi1 = 0.045 $s^{-1}$
$R^2$ = 0.95628
Δ $R^2$ = 0.02326
$k_{on}$CBD = 4.0 $s^{-1}$
$k_{off}$Chi1 = 0.044 $s^{-1}$
$R^2$ = 0.96951
Δ $R^2$ = 0.00704
$k_{on}$CBD = 8.0 $s^{-1}$
$k_{off}$Chi1 = 0.045 $s^{-1}$
$R^2$ = 0.91339
Δ $R^2$ = 0.06848
$k_{on}$CBD = 12 $s^{-1}$
$k_{off}$Chi1 = 0.045 $s^{-1}$
$R^2$ = 0.96474
Δ $R^2$ = 0.01576
$k_{on}$CBD = 16 $s^{-1}$
$k_{off}$Chi1 = 0.038 $s^{-1}$
$R^2$ = 0.9292
Δ $R^2$ = 0.0246
$k_{on}$CBD = 20 $s^{-1}$
$k_{off}$Chi1 = 0.048 $s^{-1}$
$R^2$ = 0.91493
Δ $R^2$ = 0.05093
$k_{on}$CD = 2.3 $s^{-1}$
$k_{on}$CBD = 0.23 $s^{-1}$
$k_{off}$Chi1 = 0.17 $s^{-1}$
$R^2$ = 0.88456
Δ $R^2$ = 0.10439
$k_{on}$CBD = 0.2875 $s^{-1}$
$k_{off}$Chi1 = 0.16 $s^{-1}$
$R^2$ = 0.89493
Δ $R^2$ = 0.09544
$k_{on}$CBD = 0.575 $s^{-1}$
$k_{off}$Chi1 = 0.077 $s^{-1}$
$R^2$ = 0.90004
Δ $R^2$ = 0.09386
$k_{on}$CBD = 1.15 $s^{-1}$
$k_{off}$Chi1 = 0.057 $s^{-1}$
$R^2$ = 0.97852
Δ $R^2$ = 0.01656
$k_{on}$CBD = 2.3 $s^{-1}$
$k_{off}$Chi1 = 0.049 $s^{-1}$
$R^2$ = 0.97934
Δ $R^2$ = 0.00933
$k_{on}$CBD = 4.6 $s^{-1}$
$k_{off}$Chi1 = 0.040 $s^{-1}$
$R^2$ = 0.95663
Δ $R^2$ = 0.02073
$k_{on}$CBD = 9.2 $s^{-1}$
$k_{off}$Chi1 = 0.047 $s^{-1}$
$R^2$ = 0.95247
Δ $R^2$ = 0.01811
$k_{on}$CBD = 13.8 $s^{-1}$
$k_{off}$Chi1 = 0.041 $s^{-1}$
$R^2$ = 0.91904
Δ $R^2$ = 0.05393
$k_{on}$CBD = 18.4 $s^{-1}$
$k_{off}$Chi1 = 0.039 $s^{-1}$
$R^2$ = 0.95695
Δ $R^2$ = 0.0156
$k_{on}$CBD = 23 $s^{-1}$
$k_{off}$Chi1 = 0.040 $s^{-1}$
$R^2$ = 0.93929
Δ $R^2$ = 0.0074
Events
Binding time (s)

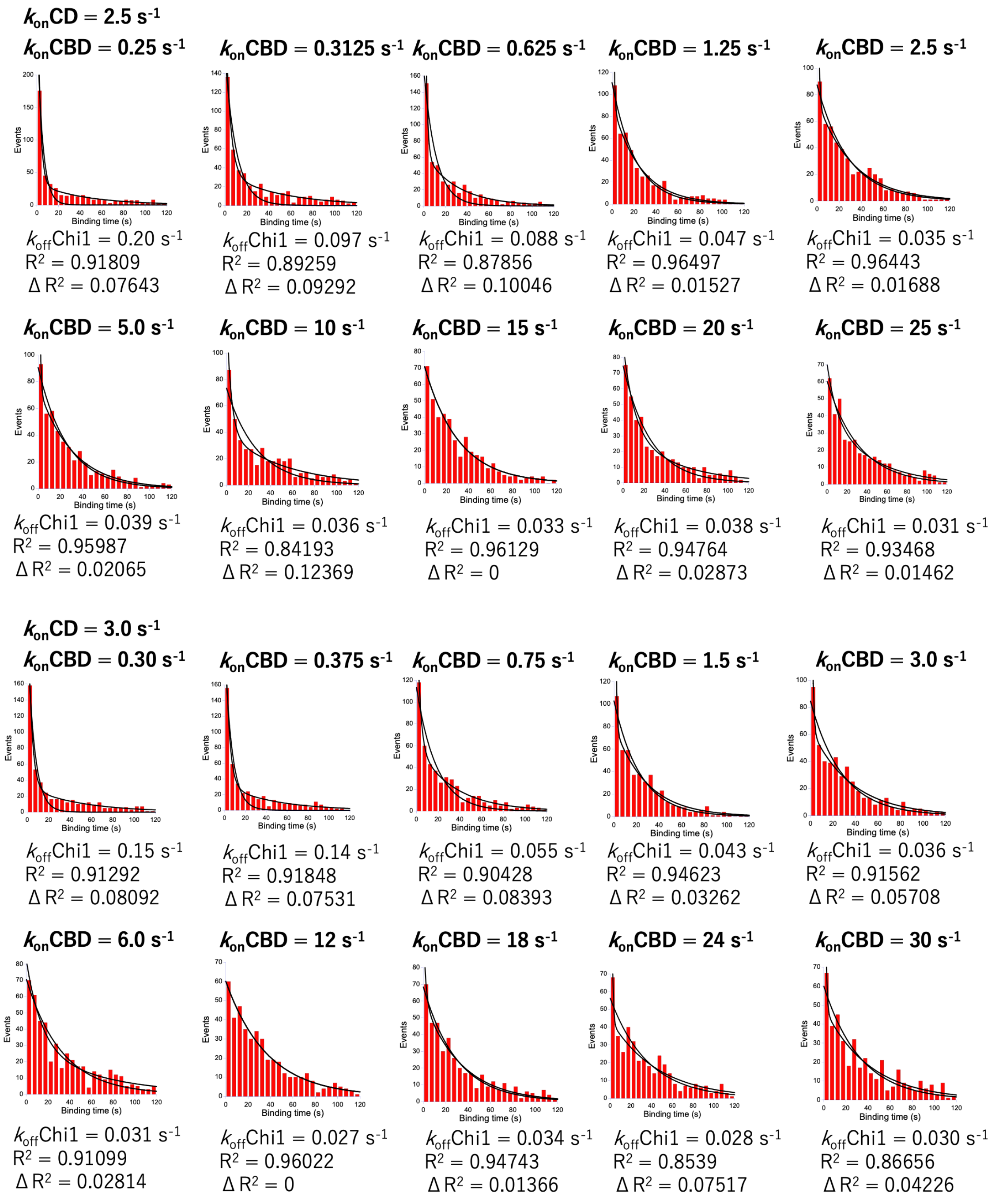

$k_{on}$CD = 2.5 $s^{-1}$
$k_{on}$CBD = 0.25 $s^{-1}$
$k_{off}$Chi1 = 0.20 $s^{-1}$
$R^2$ = 0.91809
Δ $R^2$ = 0.07643
$k_{on}$CBD = 0.3125 $s^{-1}$
$k_{off}$Chi1 = 0.097 $s^{-1}$
$R^2$ = 0.89259
Δ $R^2$ = 0.09292
$k_{on}$CBD = 0.625 $s^{-1}$
$k_{off}$Chi1 = 0.088 $s^{-1}$
$R^2$ = 0.87856
Δ $R^2$ = 0.10046
$k_{on}$CBD = 1.25 $s^{-1}$
$k_{off}$Chi1 = 0.047 $s^{-1}$
$R^2$ = 0.96497
Δ $R^2$ = 0.01527
$k_{on}$CBD = 2.5 $s^{-1}$
$k_{off}$Chi1 = 0.035 $s^{-1}$
$R^2$ = 0.96443
Δ $R^2$ = 0.01688
$k_{on}$CBD = 5.0 $s^{-1}$
$k_{off}$Chi1 = 0.039 $s^{-1}$
$R^2$ = 0.95987
Δ $R^2$ = 0.02065
$k_{on}$CBD = 10 $s^{-1}$
$k_{off}$Chi1 = 0.036 $s^{-1}$
$R^2$ = 0.84193
Δ $R^2$ = 0.12369
$k_{on}$CBD = 15 $s^{-1}$
$k_{off}$Chi1 = 0.033 $s^{-1}$
$R^2$ = 0.96129
Δ $R^2$ = 0
$k_{on}$CBD = 20 $s^{-1}$
$k_{off}$Chi1 = 0.038 $s^{-1}$
$R^2$ = 0.94764
Δ $R^2$ = 0.02873
$k_{on}$CBD = 25 $s^{-1}$
$k_{off}$Chi1 = 0.031 $s^{-1}$
$R^2$ = 0.93468
Δ $R^2$ = 0.01462
$k_{on}$CD = 3.0 $s^{-1}$
$k_{on}$CBD = 0.30 $s^{-1}$
$k_{off}$Chi1 = 0.15 $s^{-1}$
$R^2$ = 0.91292
Δ $R^2$ = 0.08092
$k_{on}$CBD = 0.375 $s^{-1}$
$k_{off}$Chi1 = 0.14 $s^{-1}$
$R^2$ = 0.91848
Δ $R^2$ = 0.07531
$k_{on}$CBD = 0.75 $s^{-1}$
$k_{off}$Chi1 = 0.055 $s^{-1}$
$R^2$ = 0.90428
Δ $R^2$ = 0.08393
$k_{on}$CBD = 1.5 $s^{-1}$
$k_{off}$Chi1 = 0.043 $s^{-1}$
$R^2$ = 0.94623
Δ $R^2$ = 0.03262
$k_{on}$CBD = 3.0 $s^{-1}$
$k_{off}$Chi1 = 0.036 $s^{-1}$
$R^2$ = 0.91562
Δ $R^2$ = 0.05708
$k_{on}$CBD = 6.0 $s^{-1}$
$k_{off}$Chi1 = 0.031 $s^{-1}$
$R^2$ = 0.91099
Δ $R^2$ = 0.02814
$k_{on}$CBD = 12 $s^{-1}$
$k_{off}$Chi1 = 0.027 $s^{-1}$
$R^2$ = 0.96022
Δ $R^2$ = 0
$k_{on}$CBD = 18 $s^{-1}$
$k_{off}$Chi1 = 0.034 $s^{-1}$
$R^2$ = 0.94743
Δ $R^2$ = 0.01366
$k_{on}$CBD = 24 $s^{-1}$
$k_{off}$Chi1 = 0.028 $s^{-1}$
$R^2$ = 0.8539
Δ $R^2$ = 0.07517
$k_{on}$CBD = 30 $s^{-1}$
$k_{off}$Chi1 = 0.030 $s^{-1}$
$R^2$ = 0.86656
Δ $R^2$ = 0.04226
Events
Binding time (s)

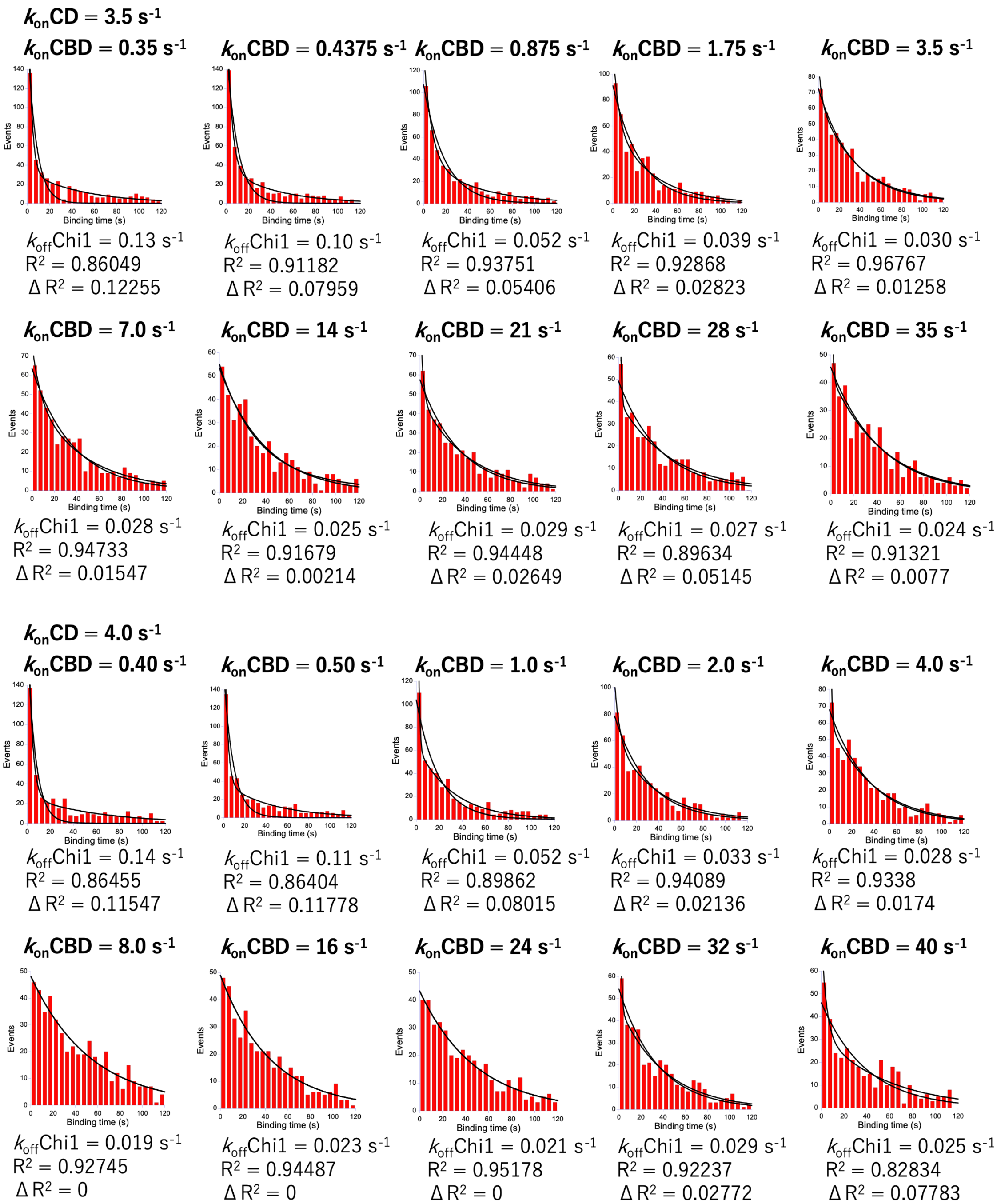

$k_{on}$CD = 3.5 $s^{-1}$
$k_{on}$CBD = 0.35 $s^{-1}$
$k_{off}$Chi1 = 0.13 $s^{-1}$
$R^2$ = 0.86049
$\Delta R^2$ = 0.12255
$k_{on}$CBD = 0.4375 $s^{-1}$
$k_{off}$Chi1 = 0.10 $s^{-1}$
$R^2$ = 0.91182
$\Delta R^2$ = 0.07959
$k_{on}$CBD = 0.875 $s^{-1}$
$k_{off}$Chi1 = 0.052 $s^{-1}$
$R^2$ = 0.93751
$\Delta R^2$ = 0.05406
$k_{on}$CBD = 1.75 $s^{-1}$
$k_{off}$Chi1 = 0.039 $s^{-1}$
$R^2$ = 0.92868
$\Delta R^2$ = 0.02823
$k_{on}$CBD = 3.5 $s^{-1}$
$k_{off}$Chi1 = 0.030 $s^{-1}$
$R^2$ = 0.96767
$\Delta R^2$ = 0.01258
$k_{on}$CBD = 7.0 $s^{-1}$
$k_{off}$Chi1 = 0.028 $s^{-1}$
$R^2$ = 0.94733
$\Delta R^2$ = 0.01547
$k_{on}$CBD = 14 $s^{-1}$
$k_{off}$Chi1 = 0.025 $s^{-1}$
$R^2$ = 0.91679
$\Delta R^2$ = 0.00214
$k_{on}$CBD = 21 $s^{-1}$
$k_{off}$Chi1 = 0.029 $s^{-1}$
$R^2$ = 0.94448
$\Delta R^2$ = 0.02649
$k_{on}$CBD = 28 $s^{-1}$
$k_{off}$Chi1 = 0.027 $s^{-1}$
$R^2$ = 0.89634
$\Delta R^2$ = 0.05145
$k_{on}$CBD = 35 $s^{-1}$
$k_{off}$Chi1 = 0.024 $s^{-1}$
$R^2$ = 0.91321
$\Delta R^2$ = 0.0077
$k_{on}$CD = 4.0 $s^{-1}$
$k_{on}$CBD = 0.40 $s^{-1}$
$k_{off}$Chi1 = 0.14 $s^{-1}$
$R^2$ = 0.86455
$\Delta R^2$ = 0.11547
$k_{on}$CBD = 0.50 $s^{-1}$
$k_{off}$Chi1 = 0.11 $s^{-1}$
$R^2$ = 0.86404
$\Delta R^2$ = 0.11778
$k_{on}$CBD = 1.0 $s^{-1}$
$k_{off}$Chi1 = 0.052 $s^{-1}$
$R^2$ = 0.89862
$\Delta R^2$ = 0.08015
$k_{on}$CBD = 2.0 $s^{-1}$
$k_{off}$Chi1 = 0.033 $s^{-1}$
$R^2$ = 0.94089
$\Delta R^2$ = 0.02136
$k_{on}$CBD = 4.0 $s^{-1}$
$k_{off}$Chi1 = 0.028 $s^{-1}$
$R^2$ = 0.9338
$\Delta R^2$ = 0.0174
$k_{on}$CBD = 8.0 $s^{-1}$
$k_{off}$Chi1 = 0.019 $s^{-1}$
$R^2$ = 0.92745
$\Delta R^2$ = 0
$k_{on}$CBD = 16 $s^{-1}$
$k_{off}$Chi1 = 0.023 $s^{-1}$
$R^2$ = 0.94487
$\Delta R^2$ = 0
$k_{on}$CBD = 24 $s^{-1}$
$k_{off}$Chi1 = 0.021 $s^{-1}$
$R^2$ = 0.95178
$\Delta R^2$ = 0
$k_{on}$CBD = 32 $s^{-1}$
$k_{off}$Chi1 = 0.029 $s^{-1}$
$R^2$ = 0.92237
$\Delta R^2$ = 0.02772
$k_{on}$CBD = 40 $s^{-1}$
$k_{off}$Chi1 = 0.025 $s^{-1}$
$R^2$ = 0.82834
$\Delta R^2$ = 0.07783
Events
Binding time (s)

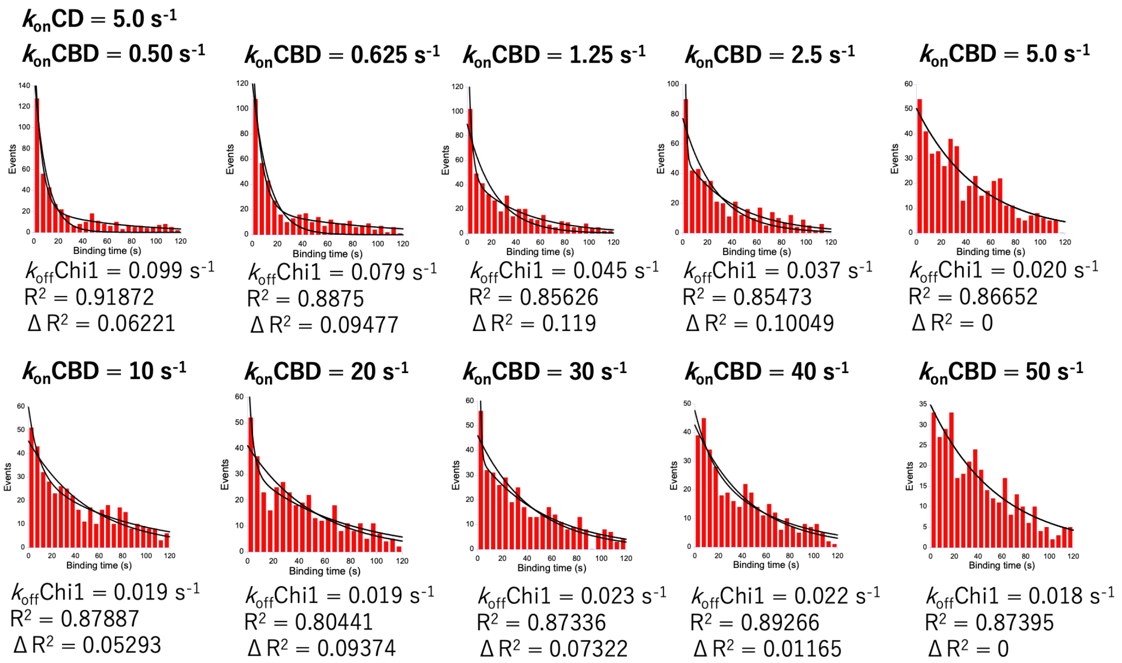


**Fig. S4 Binding time distribution simulations with different combinations of $k_{on}$ values.**

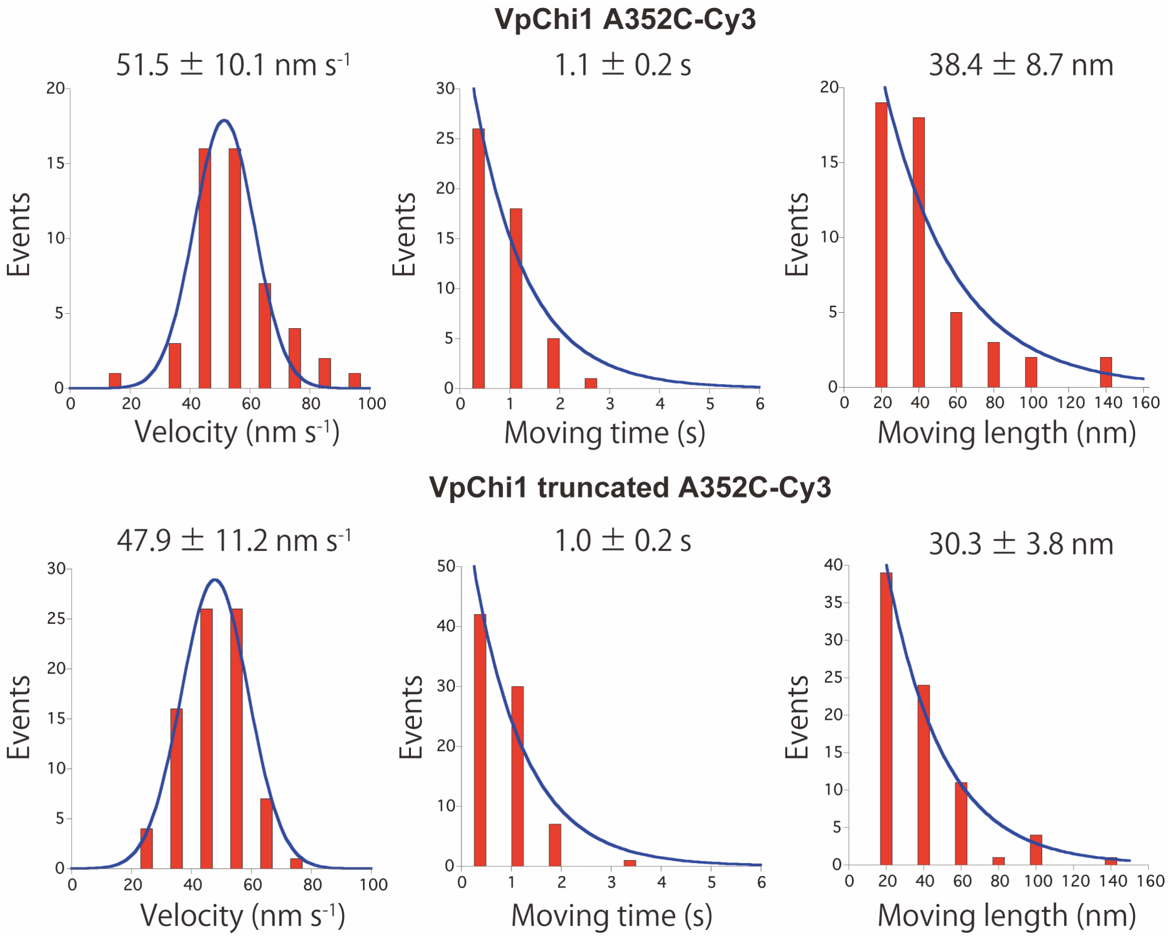


**Fig. S5 Distributions of moving velocities and moving length of VpChi1-A352C-Cy3 and the truncated mutant analyzed by HS-AFM.** Enzyme molecules moving on crystalline chitin were observed in 50 mM sodium phosphate buffer (pH 6.0) at 25°C at 5 fps. The centroids of individual molecules were tracked, and velocities were calculated from their moving lengths and times.

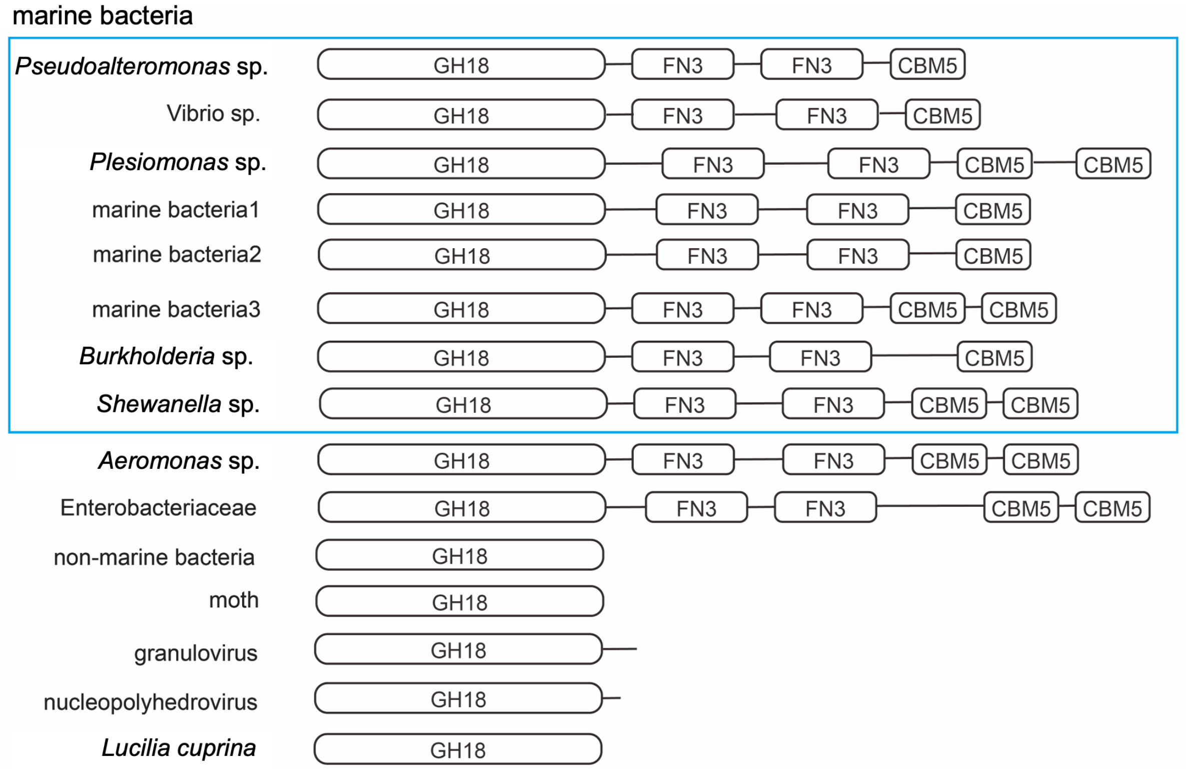


**Fig. S6 Domain compositions of the chitinases.**

```
import random
import math

k = 0.47 #koff-CBD 0.47 /s or koff-CD 0.25 /s

x = 1 #number of molecules
times=[]

while x<=500:

        t1 = 0 #binding_time s
        t1 = math.log(random.random())/-k
        times.append(t1)

        x=x+1

print(times)
```

**Fig. S7 Python script for binding time simulation of single domain**

```
import random
import math

k1 = 0.25 #CDkoff /s
k2 = 0.47 #CBDkoff /s
k3 = 2.3 # rebinding rate of CD 10 to 0.25 /s
k4 = k3*1 # rebinding rate of CBM

k3L = [0.25, 0.5, 1, 1.5, 2, 2.3, 2.5, 3, 3.5, 4, 5] #n = 0-11
k4L = [0.1, 0.125, 0.25, 0.5, 1, 2, 4, 6, 8, 10] #m = 0-10

Length = 120 #simulation time s

def bindingsim(k1,k2,k3,k4):

    bindingtime = 0
    t1sum = 0 #total time of CD s
    t1b = 0 #Binding time of CD s
    t1d = 0 #Dissociated time of CD s
    t2sum = 0 #total time of CBD s
    t2b = 0 #Binding time of CBD s
    t2d = 0 #Dissociated time of CBD s
    bindingtimes1=[] #Binding time list of CD
    dissocationtimes1=[] #Dissociated time list of CD
    bindingtimes2=[] #Binding time list of CBD
    dissocationtimes2=[] #Dissociated time list of CBD

    while t1sum < Length:

        t1b = math.log(random.random())/-k1
        bindingtimes1.append(t1b)
        t1d = math.log(random.random())/-k3
        dissocationtimes1.append(t1d)
        t1sum = t1sum + t1b + t1d

    while t2sum < Length:

        t2b = math.log(random.random())/-k2
        bindingtimes2.append(t2b)
        t2d = math.log(random.random())/-k4
        dissocationtimes2.append(t2d)
        t2sum = t2sum + t2b + t2d

    L1b = len(bindingtimes1)
    L2b = len(bindingtimes2)
    L1d = len(dissocationtimes1)
    L2d = len(dissocationtimes2)
    n1 = 0
    n2 = 0
    tcheck = 0

    if bindingtimes2[0] <= bindingtimes1[0]:
        tcheck = bindingtimes1[0]+dissocationtimes1[0]
        while n2+1 < L2b:
            while tcheck - bindingtimes2[n2] >= 0:
                if tcheck - dissocationtimes1[n1] - bindingtimes2[n2] <= 0:
                    if n2 >= 1:
                        bindingtime = sum(bindingtimes2[0:n2])+sum(dissocationtimes2[0:n2-1])

                        Binding_times.append(bindingtime)
                        return(Binding_times)
                    else:
                        bindingtime = bindingtimes2[0]
                        Binding_times.append(bindingtime)
                        return(Binding_times)

                else:
                    tcheck = tcheck - bindingtimes2[n2] - dissocationtimes2[n2]

                    n2 = n2+1
                    if n2 == L2b:
                        bindingtime = Length
                        Binding_times.append(bindingtime)
                        return(Binding_times)
            if n1+1 < L1b:
```

```
                                        n1 = n1+1
                                        tcheck = tcheck + bindingtimes1[n1]+dissocationtimes1[n1]

        elif bindingtimes2[0] > bindingtimes1[0]:
                tcheck = bindingtimes2[0]+dissocationtimes2[0]
                while n1+1 < L1b:
                        while tcheck - bindingtimes1[n1] >= 0:
                                if tcheck - dissocationtimes2[n2] - bindingtimes1[n1] <= 0:
                                        if n1 >= 1:
                                                bindingtime                                        =
sum(bindingtimes1[0:n1])+sum(dissocationtimes1[0:n1-1])
                                                Binding_times.append(bindingtime)
                                                return(Binding_times)
                                        else:
                                                bindingtime = bindingtimes1[0]
                                                Binding_times.append(bindingtime)
                                                return(Binding_times)

                                else:
                                        tcheck     =     tcheck     -     bindingtimes1[n1]     -
dissocationtimes1[n1]
                                        n1 = n1+1
                                        if n1 == L1b:
                                                bindingtime = Length
                                                Binding_times.append(bindingtime)
                                                return(Binding_times)
                        if n2+1 < L2b:
                                n2 = n2+1
                                tcheck = tcheck + bindingtimes2[n2]+dissocationtimes2[n2]

n = 0

while n < 11:
        k3 = k3L[n]
        m = 0
        while m < 10:
                k4 = k3*k4L[m]

                x = 1 #counts
                Binding_times=[]

                while x<500:
                        bindingsim(k1,k2,k3,k4)
                        x=len(Binding_times)
                k4f = '{:.3f}'.format(k4)
                filename = 'Vp_kon3_'+str(k3)+'_kon4_'+str(k4f)
                f = open(filename+'.txt', mode='w')
                print(Binding_times, file=f)
                f.close()

                m = m+1

        n = n+1
```

**Fig. S8 Python script for binding time simulation of double domain**

**Legends for Supplementary Movies**

**Supplementary movie 1**

**Binding and dissociation events of VpChi1 molecules.**

VpChi1-Cy3 (25 pM) in 50 mM sodium phosphate buffer (pH 6.0) was applied to the chitin-coated glass surface and imaged at 4 fps using a laser power density of 0.14 μW $\mu m^{-2}$. The movie is played at 10× real-time speed.

**Supplementary movie 2**

**Binding and dissociation events of VpChi1-truncated molecules.**

VpChi1-truncated-Cy3 (25 pM) in 50 mM sodium phosphate buffer (pH 6.0) was applied to the chitin-coated glass surface and imaged at 4 fps using a laser power density of 0.14 μW $\mu m^{-2}$. The movie is played at 10× real-time speed.

**Supplementary movie 3**

**Binding and dissociation events of VpChi1-2ndCBD molecules.**

VpChi1-2ndCBD-Cy3 (40 pM) in 50 mM sodium phosphate buffer (pH 6.0) was applied to the chitin-coated glass surface and imaged at 10 fps using a laser power density of 0.28 μW $\mu m^{-2}$. The movie is played at 10× real-time speed.

**Supplementary movie 4**

**Movement of VpChi1 molecules visualized by HS-AFM.**

VpChi1 was imaged in 50 mM sodium phosphate buffer (pH 6.0) at 25 °C and 5 fps using high-speed atomic force microscopy. The movie is played at 3× real-time speed.

**Supplementary movie 5**

**Movement of VpChi1-truncated molecules visualized by HS-AFM.**

VpChi1-truncated was imaged in 50 mM sodium phosphate buffer (pH 6.0) at 25 °C and 5 fps using high-speed atomic force microscopy. The movie is played at 3× real-time speed.
